# PRDM16 is necessary for sensory neuronal development in the Trigeminal Ganglion

**DOI:** 10.64898/2026.02.01.702826

**Authors:** Fahmida Raha, Qiman Gao, Lomeli C. Shull, Kristin B. Artinger

## Abstract

**Background:** Cranial neural crest cells (cNCC) generate craniofacial cartilage, bone, and peripheral neurons and glia, and birth defects arise when the cartilage/neuronal/glial progenitor fail to differentiate. PRDM16 is a transcriptional regulator containing both zinc-finger and SET domains, implicated in craniofacial development and orofacial clefting, but its role in cranial sensory ganglion formation has not been defined.

**Results:** Here, we demonstrate that *prdm16* is required for trigeminal ganglion (TG) assembly and sensory neurogenesis from cranial neural crest lineages. In zebrafish, *prdm16* is expressed in TG beginning by 18 hours post fertilization (hpf) and persists through the later developmental stage at 48 hpf. In the *prdm16* loss-of-function zebrafish, fewer HuC+ TG neurons are present at 24 hpf and 48 hpf, along with reduced overall ganglion size. Live imaging in Tg(*sox10*:mRFP; *elavl3*:GFP) embryos demonstrates similar numbers of *sox10*+ cNCCs migrating to the TG region and reduced cell numbers and overall smaller size of TG in *prdm16-/-*. Acetylated β-tubulin immunostaining shows fewer trigeminal axon projections early and an altered projection pattern by 48 hpf. A reduction in a defined sensory neuron population, *p2rx3b*+ cells displayed a weaker signal and decreased cell number in *prdm16-/-* TG. Transcriptomic analysis of FACS-isolated *sox10*+ cranial neural crest cells supported reduced expression of key neurogenic and sensory lineage genes. Finally, in mouse embryos, PRDM16 is expressed in TG neurons, and *Prdm16^csp1/csp1^* embryos exhibited reduced TG volume, area and fewer HuC+ neurons at E18.5.

**Conclusion:** Together, these data identify Prdm16 as a conserved regulator of trigeminal ganglion growth and sensory neuron differentiation, linking PRDM-family chromatin regulators to the development of the peripheral sensory nervous system.

## Introduction

Most of the primary sensory organs are derived from two transient embryonic stem cell populations: the neural crest and the cranial placodes ^1,2^. A coordinated sequence of induction, migration, and differentiation organizes the precursors into groups of cells that condense into ganglia and extend axons. Cranial neural crest cells (cNCCs) give rise to a remarkable range of differentiated cell types through a highly conserved lineage progression, in which multipotent cells transition into more fate-restricted progenitors. This includes more restricted progenitor cells that gives rise to craniofacial tissues such as bone, cartilage, and connective tissue ^3,4^, and also contribute to the peripheral nervous system, including sensory and autonomic ganglia and their associated glia ^5–9^.

Neural crest development and differentiation into sensory neurons is controlled by tightly coordinated gene regulatory networks (GRNs) that govern induction, specification, migration, survival, and terminal differentiation^10–12^. This GRN activates tissue-specific transcriptional programs to generate distinct cranial neural crest lineages, including a cartilage/neuron/glial progenitor that eventually differentiates into craniofacial cartilage and connective tissues as well as peripheral glia and sensory neurons^4,13^. Increasing evidence indicates that this regulation is not only controlled by transcription factor networks but also by chromatin modifiers that have important regulatory roles in cNCC and craniofacial development ^14,15^. However, not much is known about the epigenetic factors important for trigeminal ganglion development.

Migrating neural crest cells (NCCs) reach prospective ganglion sites first, but their differentiation is delayed until placodal cells have migrated and begun differentiating^16,17^. Interactions between NCCs and placode cells then give rise to craniosensory ganglia, including the largest and first one to be formed, the trigeminal ganglion (TG)^17–19^. In zebrafish, the neural crest is specified as early as 11 hours post fertilization (hpf) at the lateral edges of the anterior neural plate border, and they begin to migrate away from the neural ectoderm around 14 hpf ^20^. After reaching the proper destination, these NCCs collaborate with placodal cells in the formation of the cranial ganglia. The cNCCs begin migration as a single mass and subsequently branch into discrete streams, three of which are broadly conserved across vertebrates: trigeminal, hyoid, and post-otic^21,22^. The trigeminal stream arises from the midbrain and rhombomeres 1 and 2 of the hindbrain, contributes to neurons within the trigeminal ganglion, and populates the orofacial prominences and mandibular arch that form the skeletal elements of the upper and lower jaw^23,24^. The TG contains the cell bodies of sensory neurons that convey touch, pain, temperature, and proprioceptive information from the face and oral region to the brain via the trigeminal nerve (cranial nerve V). Within the ganglion, sensory neuron subtypes are distinguished in part by the expression of specific receptors ^25,26^. Mechanosensory neurons express mechanosensory channels, including PIEZO2^27^, and are often associated with NTRK2 (TRKB) and NTRK3 (TRKC) sensory lineages^7,28^. Heat and irritant-responsive populations frequently express Transient Receptor Potential (TRP) channels that function in sensory perception, including TRPV1 and TRPA1^29,30^, while many nociceptors are required for pain sensation combine NTRK1 (TRKA)-linked trophic signaling with ATP sensing through the purinergic receptor P2X3^31–33^. P2X3 (encoded by *P2RX3*) is an extracellular ATP–gated cation channel that is enriched in primary nociceptive sensory neurons, including trigeminal ganglion neurons, and links ATP released during tissue stress or inflammation to neuron depolarization^31,34,35^. P2X3 purinergic receptor has also been shown to promote facial pain by activating neurons in TG ^36,37^. With subtype specification, TG neurons must also extend axons to appropriate peripheral targets in the face and in the brainstem to establish functional sensory circuits ^38^. Developmental studies show that trigeminal axon branching and targeting can be halted or changed due to multiple factors, including axon guidance cues and genes functioning in proper TG formation^39–41^. Inhibition of axon outgrowth can also be seen by repressing signals from particular sensory neuron subtypes^42^.

The PRDM (PRDI-BF1 and RIZ homology domain containing) protein family acts as tissue-specific transcription factors and are expressed in neural crest-derived tissues including the head folds in the early neural plate, pharyngeal arches, and the peripheral nervous system ^43,44^. PRDM proteins are notable for their ability to bind DNA directly through zinc-finger domains, to influence both repressive H3K9me3 and active H3K4me3 histone marks via a PR/SET methyltransferase domain, and to engage chromatin regulators through protein–protein interactions ^43,45–47^. Several PRDM factors have been implicated in neurocristopathies; notably, *PRDM16* has been linked to human cleft lip and palate ^48–50^ and mutations in mice result in cleft palate and a persistent open-mouth phenotype ^48,49^. In zebrafish, *prdm16* loss-of-function causes hypoplastic or missing posterior viscerocranium cartilage, accompanied by defects in chondrocyte polarity and a reduced, clefted neurocranium ^51,52^. However, despite its established roles in craniofacial development and broader expression in the head folds and central nervous system, the role of *prdm16* in cNCC differentiation into neuronal fates has not been examined. PRDM16 has been linked to migraine pain^53,54^, raising the possibility that PRDM16 influences sensory neuron development and function in the TG but remains largely unexplored.

Here, we test the requirement for *prdm16* in TG formation, neurogenesis and axon guidance from cNCC to the TG. We show that *prdm16* is expressed in the developing TG in both neural crest and placodal cells. In both zebrafish and mouse *prdm16* loss-of-function models, the TG is smaller and contains fewer neurons, indicating a conserved requirement for *prdm16* in ganglion growth. We further find defects in trigeminal axon projection patterns and a reduction in a defined sensory neuron population marked by *p2rx3b*-sensory neurons. Finally, transcriptomic data from isolated cNCCs support a decrease in the neurogenesis transcription program in *prdm16* mutants. Together, these data suggest the role for *prdm16*, a chromatin-associated transcriptional regulator, in coordinating TG formation and sensory neuron differentiation.

## Results

### *prdm16* is expressed in both neural crest and placode-derived trigeminal neurons during development and maturation

To define the spatial and temporal pattern of *prdm16* expression in zebrafish relative to TG formation, we performed in situ Hybridization Chain Reaction (HCR) in situ hybridization using *prdm16* and the neuronal cell body marker *elavl3* across a developmental time course (14, 18, 24, and 48 hpf. *elavl3* (HuC) is expressed in neuronal cell bodies in the TG (marked in white dotted circles in Figure 1) and we examined the proximity of *prdm16* expression to *elavl3* (HuC) (Figure 1). At 14 hpf (Figure 1A-A’’), at the time that NCCs begin to migrate from the neural plate border, *prdm16* expression was not observed in proximity to the developing TG. By 18 hpf (Figure 1B-B’’), *prdm16* begins to be expressed adjacent to the TG, and this TG-associated expression is clearly detectable at 24 hpf (Figure 1C-C’’). It remains expressed at 48 hpf, when the ganglion is more mature, primarily in the ventral domain of *elavl3* expression (Figure 1D-D’’). Across later stages, *prdm16* signal was also visible in additional cranial domains, including the hindbrain and pharyngeal arch regions, consistent with broader expression during craniofacial development (Figure 1D-D’’). Because TG assembly depends on the interaction of neural crest and placode-derived cells, we also examined the expression *prdm16* in each lineage type (Figure 1E-E’’ and Supplementary Figure 1). In transgenic embryos with the neural crest^55,56^ (Tg:*sox10:e*GFP), HCR of *prdm16* shows overlapping expression with *sox10+* cells in the vicinity of the TG at 48 hpf (Figure 1E-E’’). *prdm16* is also detected in the placode expressing domain of *six1b*+ cells^57^ (placode marker) in the TG region (Supplementary Figure 1), indicating that *prdm16* is present in both neural crest and placode-derived trigeminal cell populations at 48 hpf. Together, these data suggest the presence of *prdm16* in the right place and time to influence TG morphogenesis.

**Figure 1.**
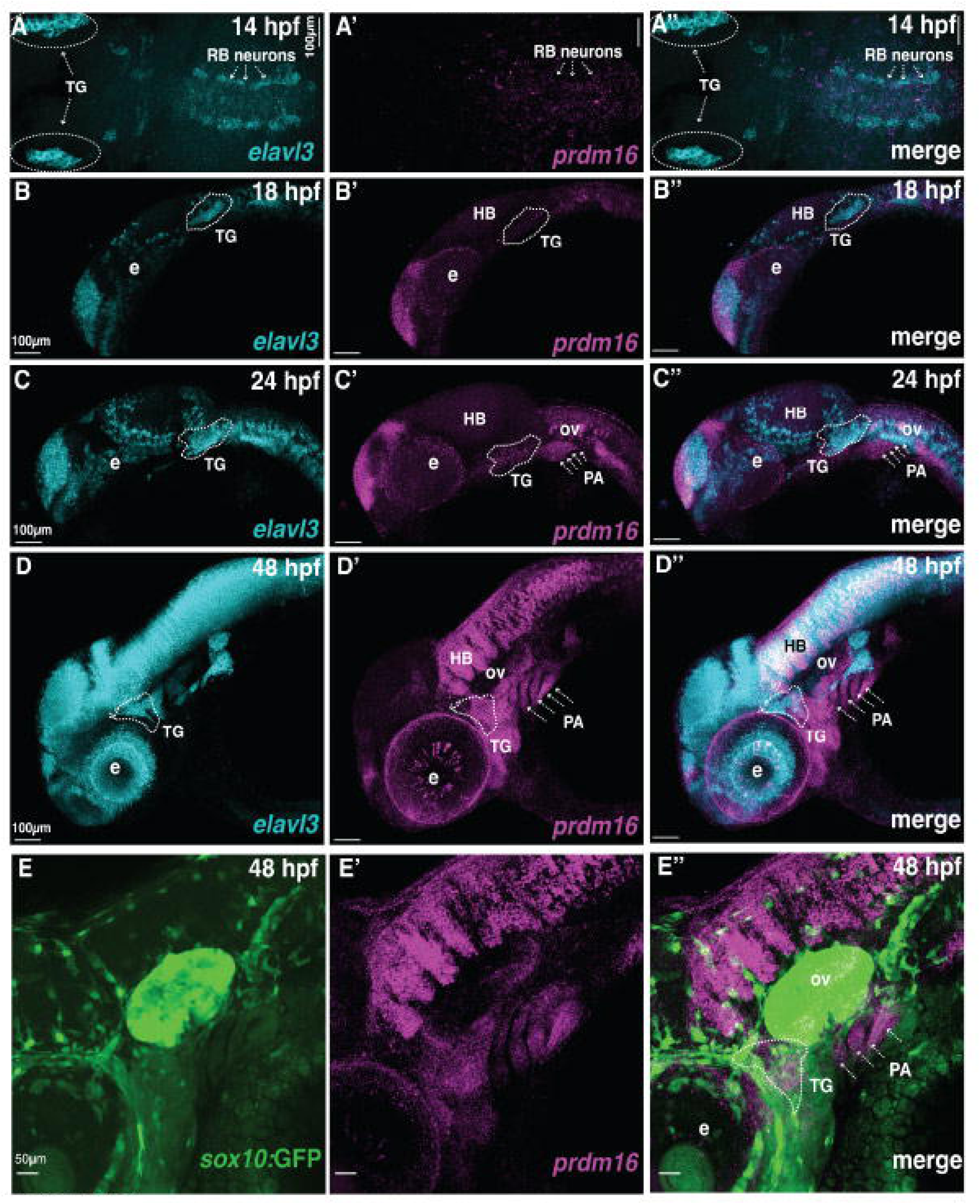
*prdm16* expression is present in the region of the TG by 18 hpf and persists throughout ganglion maturation. Whole-mount HCR in situ hybridization was used to localize *prdm16* transcripts relative to the developing trigeminal ganglion (TG) neuronal marker *elavl3* (HuC) across early stages of TG assembly. (A–D) Representative confocal images show *prdm16* (magenta) and *elavl3* (cyan) signal at 14 hpf (A), 18 hpf (B), 24 hpf (C), and 48 hpf (D), with corresponding merged views; white dotted outlines indicate the TG region. Dorsal view, anterior to the left (A-A’’) at 14 hpf, *prdm16* signal is detected in Rohan-Beard (RB) neurons, but not adjacent to the forming TG. (B-E) Lateral view, anterior to the left. (B-B’’) By 18 hpf, *prdm16* expression becomes TG-proximal and remains readily detectable at (C-C’’) 24 hpf and (D-D’’) 48 hpf, when additional cranial expression domains are also apparent (including hindbrain and pharyngeal arches). (E) At 48 hpf, *prdm16* HCR signal overlaps with *sox10*:GFP-positive neural crest cells in the TG vicinity, supporting *prdm16* expression within neural crest–associated trigeminal cell populations. Scale bars: 100 µm (A–D) and 50 µm (E). . Abbreviations: TG, trigeminal ganglion; HB, hindbrain; PA, pharyngeal arches; RB neurons, Rohon–Beard neurons; ov, otic vesicle; e, eye; hpf, hours post fertilization.

### *prdm16* mutants have a persistent reduction in trigeminal ganglion neuron cell number and volume

Previously, we generated several *prdm16* mutant alleles by Crispr/Cas9 mutagenesis, all of which is predicted to have a frameshift mutation that disrupts the coding sequence upstream of the PR/SET domain responsible for functional histone methyltransferase activity^51^. For this study, we used the *prdm16*^CO1027^ allele. Loss of *prdm16* causes moderate overall craniofacial phenotypes, namely mild hypoplasia of the cartilage in the developing zebrafish pharyngeal skeleton. However, the role of *prdm16* in the formation of peripheral sensory neurons is not known.

To determine whether *prdm16* is required for proper TG development, we examined and quantified TG neurons in *prdm16*-/- embryos across developmental stages of ganglion development. Since *prdm16* is strongly expressed near the TG around 24 hpf, we examined neuronal differentiation at 24 hpf and at 48 hpf when the TG is more mature. At 24 hpf (1 day post fertilization, dpf), HuC immunolabeling revealed a compact cluster of differentiated neurons in the outlined TG region in wildtype embryos (Figure 2A–A’’). In contrast, *prdm16*-/- embryos displayed a smaller HuC+ domain within the TG territory at the same stage (Figure 2B–B’’). 3D-quantification confirmed a significant decrease in HuC+ TG neuron number in *prdm16*-/- compared to wildtype at 24 hpf (Figure 2C, n=7 per group).

**Figure 2.**
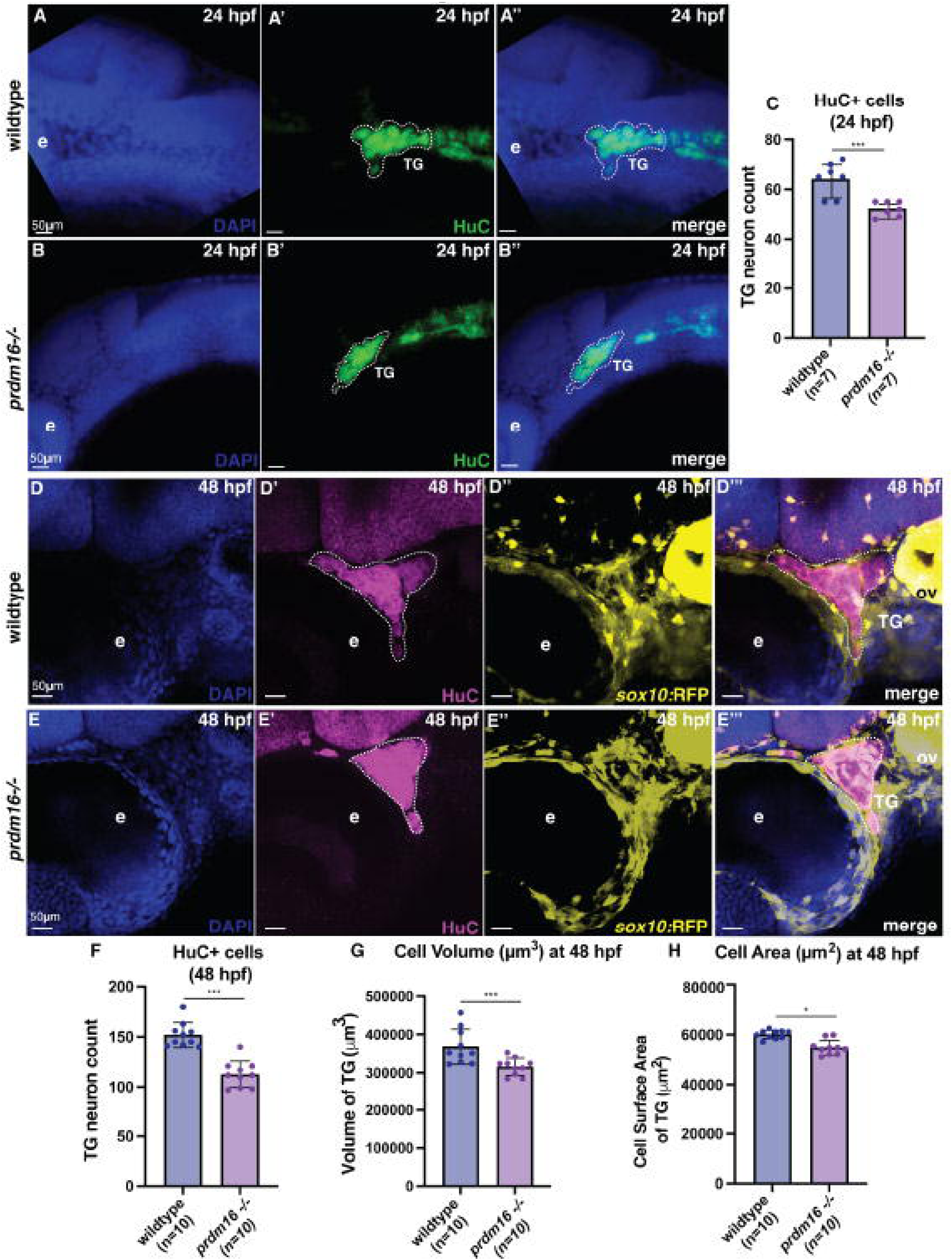
*prdm16* loss shows a reduction in TG neuron number and ganglion size. Whole-mount immunofluorescence shows fewer differentiated trigeminal neurons and reduced trigeminal ganglion (TG) morphometrics in *prdm16*-/- embryos compared with wildtype at 24 and 48 hpf. Lateral view, anterior to the left. (A–B) Representative lateral views at 24 hpf stained for DAPI (blue) and HuC (green), and merge of these two channels. The TG regions are outlined with white dashed lines. (A–A’’) Wildtype embryos show a compact HuC+ TG cluster. (B–B’’) *prdm16-/-* embryos show a visibly smaller HuC+ TG domain at the same stage. (C) Quantification of HuC+ TG neurons at 1 dpf (24 hpf) reveals a significant reduction in *prdm16-/-* compared to wildtype (n=7 embryos per genotype, p < 0.001, ***). (D–E) Representative lateral views at 48 hpf stained for DAPI (blue), HuC (magenta), and *sox10*:RFP (yellow), with the TG outlined (white dashed line). (D-D’’) Wildtype embryos show a well-defined HuC+ TG associated with *sox10*:RFP+ neural crest–derived cells. (E–E’’) *prdm16-/-* embryos retain *sox10*:RFP+ cells in the region but display reduced HuC signal and a smaller TG outline. (F) Quantification of HuC+ TG neuron number at 48 hpf (2 dpf) confirms a persistent and significant reduction in *prdm16-/-* embryos (n=10 per genotype; p < 0.001, ***). (G–H) 3D morphometric measurements at 48 hpf (2 dpf) show decreased TG volume (G) and reduced TG surface area (H) in *prdm16−/−* embryos relative to wildtype (n=10 per genotype; p < 0.001, ***; p < 0.05, *). Scale bars: 50 µm. Abbreviations: TG, trigeminal ganglion; ov, otic vesicle; e, eye; hpf, hours post fertilization; dpf, days post fertilization.

This neuronal deficit persists as the TG matures. At 48 hpf (2 dpf), wildtype embryos displayed a well-defined TG with strong HuC+ neuronal labeling in *sox10:*RFP+ neural crest–derived cells (Figure 2D–D’’’), whereas *prdm16*-/- embryos exhibited a smaller TG volume with reduced HuC signal despite the continued presence of *sox10:*RFP+ cells (Figure 2E–E’’’). There are some variations in the phenotype (Supplemental Figure 2), however 3D-quantification again confirmed a persistent reduction in TG neuron number at 48 hpf (Figure 2F, n=10 per group). In addition, 3D morphometric analysis revealed that *prdm16*-/- had significantly decreased TG volume (Figure 2G) and moderately decreased TG surface area (Figure 2H, n=11/15) at 48 hpf compared to wildtype. These data indicate that *prdm16* mutants have reduced overall ganglion size, accompanying the neuronal cell loss. Despite this structure variation, quantification of the TG neurons and volume in wildtype vs *prdm16* mutants, show that *prdm16*-/- embryos have a significant reduction in neuron cell number, area, and volume of the TG compared to wildtype embryos (Figure 2F-H).

### Similar numbers of neural crest cells migrate to TG despite reduced ganglion size in *prdm16* mutants

To test whether the reduced TG size in *prdm16-/-* embryos (Figure 2) indicates impaired neural crest migration or is a later differentiation defect in ganglion maturation, we live-imaged using double-transgenic embryos expressing *sox10:RFP* (which labels neural crest cells) and *elavl3:*GFP (labeling differentiating neurons). We crossed Tg(*sox10*:mRFP) and Tg(*elavl3*:GFP) lines to generate a Tg(*sox10*:mRFP;*elavl3*:GFP) line enabling simultaneously visualization of *sox10*+ cNCCs cells migrating in the TG area and differentiated TG neurons labeled by *elavl3* expression. We began imaging at 22 hpf (0 hr) when the TG is starting to coalesce and followed embryos for 13 hours (to approximately 35 hpf) (Supplemental Video 1 represents wildtype time-lapse and Supplemental Video 2 represents *prdm16-/-*). Across the time series, the overall migration path of *sox10*:RFP+ neural crest cells and accumulation at the TG site appeared comparable between wildtype and *prdm16-/-* embryos (Figure 3A–H, Supplemental Video 1-2). To quantify this, we counted *sox10*+ cells in the TG region across z-stacks and found a similar number of cells in wildtype and *prdm16-/-* embryos (Figure 3I, n=2). Together, these data indicate that loss of *prdm16* does not affect neural crest migration to the TG region during development. Despite the preserved number of neural crest cells during migration, the TG appeared smaller in *prdm16-/-*embryos throughout the time-lapse series (Supplemental Video 2). We also quantified in 2D the TG surface area from the outlined snapshots and found that the TG area was consistently reduced in mutants compared to wildtype (Figure 3J, n=2). These data suggest that the reduced TG size in *prdm16* mutants is not due to impaired neural crest cell number migrating to the TG, but rather may be a defect in later steps that drive ganglion differentiation and maturation.

**Figure 3:**
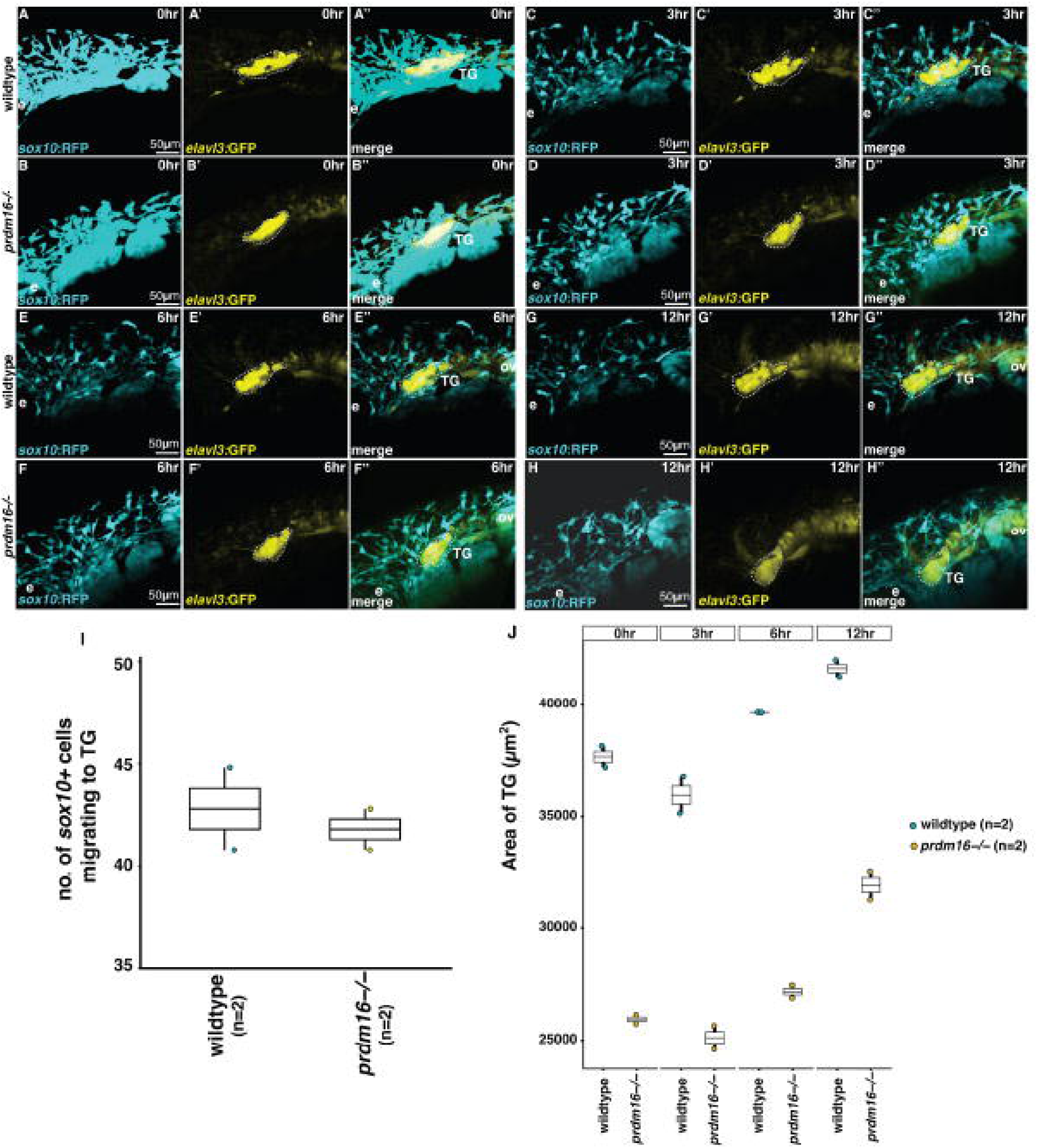
Neural crest cells migrate normally to trigeminal ganglion in *prdm16* mutants, but the ganglion is smaller during coalescence. (A–H) Still frames of lateral views with anterior to the left from live confocal time-lapse imaging of Tg(*sox10*:RFP; *elavl3*:GFP) embryos showing *sox10*:RFP making *sox10*+ cranial neural crest cells (cyan) and differentiating trigeminal neurons marked by *elavl3*:GFP (yellow) in wildtype and *prdm16-/-* embryos during TG assembly. Imaging began at ∼22 hpf (0 hr) and continued for ∼13 h (to ∼35 hpf). Representative time points are shown at 0 hr or ∼22 hpf old embryo (A-A’’ represent wildtype, B-B’’ represent *prdm16-/-*), 3 hr or ∼25 hr old embryo (C-C’’ represent wildtype, D-D’’ represent *prdm16-/-*), 6 hr or ∼28 hpf old (E-E’’ represent wildtype, F-F’’ represent *prdm16-/-*), and 12 hr or ∼34 hpf old (G-G’’ represent wildtype, H-H’’ represent *prdm16-/-).* The trigeminal ganglion region is outlined with a white dashed line in each panel. Overall, *sox10*+ migration paths and accumulation in the TG territory appear comparable between genotypes, while the outlined TG domain is visibly reduced in *prdm16*−/− across the series (see also Supplemental Videos 1–2). (I) Quantification of *sox10*+ cell number within the TG region across z-stacks shows no difference between wildtype and *prdm16*-/- embryos (n=2 per group). (J) Quantification of 2D TG surface area measured from the outlined region at matched time points shows a consistently smaller TG area in *prdm16*-/- compared with wildtype over the course of imaging (n=2 per group). Scale bar: 50 µm. Abbreviations: TG, trigeminal ganglion; ov, otic vesicle; e, eye; hpf, hours post fertilization.

To confirm that *prdm16* is acting through the altered differentiation mechanism, we also examined cell proliferation and cell death in TG and TG-proximal area (pharyngeal arches, and area between eye and otic vesicle where TG appears, Supplementary Figure 3) at the onset of TG coalescence, 24 hpf, when *sox10* cells destined to be neurons finish migrating to the TG area. We immunostained zebrafish embryos using anti phosphohistone-H3 (pHH3, cell proliferation), and anti-cleaved Casp3 (cleaved caspase-3, cell death) at 24 hpf. We counted cells expressing pHH3 (Supplementary Figure 3A-B) and cleaved Caspase-3 (Supplementary Figure 3C-D) in TG and TG-proximal area marked in white dotted circles. This broad region was chosen to consider the potential for disaggregated cells. Consistent with previous results, comparisons between wildtype and *prdm16-/-* embryos showed no significant differences in cell proliferation or cell death in either the trigeminal or trigeminal proximal area, where NCCs migrate in the same stream at 24 hpf (n=4, ns, Supplementary Figure 3 E-F). Together these results provide evidence that *prdm16* loss does not affect neural crest migration, proliferation, or apoptosis of cNCC derived TG but rather *prdm16* acts tissue-specifically in the TG and TG-proximal area for maintaining proper cell fate differentiation of neural crest to TG neurons.

### *prdm16* loss reduces the number of trigeminal axon projections and alters projection morphology

Next, we investigated whether *prdm16* is required for trigeminal peripheral nerve outgrowth and innervation by examining axonal projections. We immunostained embryos with an antibody against acetylated β-tubulin, which labels nerve fibers ^58,59^, including peripheral nerve fibers in zebrafish, and allows trigeminal branches to be readily visualized.

At 24 hpf (1 dpf), wildtype embryos displayed the typical array of fine acetylated tubulin processes. Several distinct axon projections could be seen radiating away from the ganglion into the surrounding craniofacial region (Figure 4A-A’’). In *prdm16* embryos, TG-associated fibers were present, but the pattern of cranial innervation was frequently disturbed. Some branches of the TG peripheral nerves were missing or shortened. The remaining nerves often (in 8 out of 11 mutant embryos) showed thicker processes close to the TG itself (Figure 4B-B’’). Quantification confirmed a significant reduction in the number of TG axon projections in mutants compared with wildtype at 24 hpf (Figure 4C, n=8 per group).

**Figure 4.**
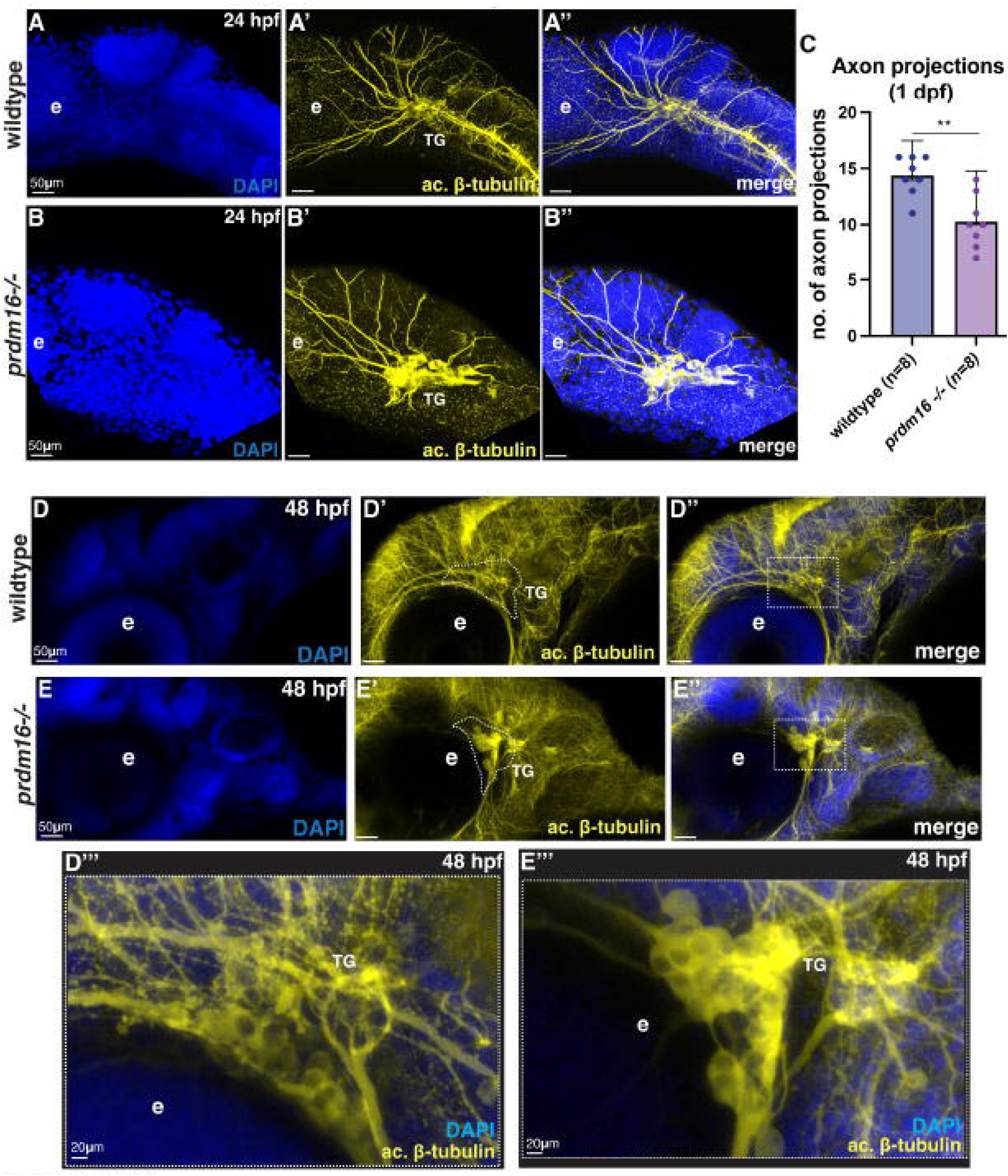
*prdm16* is required for normal trigeminal axon projections and branching morphology. Whole-mount immunofluorescence for acetylated β-tubulin was used to visualize trigeminal nerve fibers in wildtype and *prdm16*-/- embryos at 24 hpf and 48 hpf. Lateral view, anterior to the left. (A-B) At 24 hpf, representative lateral views show DAPI (blue), acetylated β-tubulin (yellow), and merged images. (A-A’’) Wildtype embryos display multiple stereotyped TG-associated projections extending into the craniofacial region, whereas (B-B’’) *prdm16*-/- embryos frequently show missing or shortened branches and thicker fibers near the TG at 24 hpf. (C) Quantification of the number of trigeminal axon projections at 1 dpf (wildtype n=8; *prdm16*-/- n=8; p < 0.01, **) indicates statistical significance. (D–E) At 48 hpf, acetylated β-tubulin labeling reveals a dense and finely branched trigeminal projection network in wildtype (D-D’’), while *prdm16*-/- embryos show reduced fine branching and abnormal fiber accumulations near the TG (E-E’’). (D’’’-E’’’) Higher-magnification views of boxed regions in (D’’) and (E’’), highlighting local branching architecture around the TG. Statistical test used was the Kruskal-Wallis non-parametric ANOVA test. Scale bars: 50 µm (A–B’’, D–E’’), 20 µm (D’’’–E’’’). Abbreviations: TG, trigeminal ganglion; e, eye; hpf, hours post fertilization; dpf, days post fertilization.

By 48 hpf (2 dpf), wildtype embryos showed a dense network of cranial fibers surrounding the eye and dense wrapping around and traversing the TG neuronal cell body territory (Figure 4D-D’’). High-magnification views showed many thin branches and a dense local projection pattern in wildtype (Figure 4D’’’) that are missing in *prdm16* mutants (Figure 4E’’’). In contrast, *prdm16-/-* embryos exhibited reduced fine branching and irregular acetylated tubulin–positive accumulation surrounding the TG (Figure 4E’’’). Together, these data suggest that *prdm16* is required for trigeminal axons to develop a normal branching pattern and proper innervation.

### *prdm16* is required for *p2rx3b*-type sensory neuron differentiation in the trigeminal ganglion

The trigeminal ganglion contains multiple sensory neuron classes, including mechanosensory and nociceptive-like populations, distinguished by their receptor expression patterns^60,61^. To further examine the aspects of sensory neuron differentiation, we investigated a particular subtype of neurons expressing *p2rx3b*, which encodes a P2X3 purinergic receptor subunit expressed in zebrafish trigeminal neurons^32^. P2X3 channels are extracellular ATP–gated cation channels that are associated with primary sensory neurons and nociceptive signaling^33^. P2X3 has also been studied in sensory neurons as key ATP-sensitive receptors involved in pain transduction and sensitization ^34^. *p2rx3b* is expressed in a high number of zebrafish trigeminal ganglia neurons starting from 24 hpf through 72 hpf^62,63^, and is functionally important, which makes it a good candidate to study the effect of *prdm16* knockout on a subpopulation of sensory neurons in TG.

We performed HCR in situ hybridization to study *p2rx3b+* trigeminal sensory neurons differentiation in *prdm16* mutants. At 48 hpf, in *prdm16-/-* embryos, *p2rx3b* signal was still present in the TG but appeared reduced in *sox10:*RFP+ cells (Figure 5B-B’’’) compared to wildtype embryos (Figure 5A-A’’’). Consistent with this, quantification showed a significant decrease in the number of *p2rx3b+* sensory neurons in *prdm16* mutants compared to wildtype at 48 hpf (Figure 5C, wildtype n=7; *prdm16-/-*, n=6). We also measured *p2rx3b* HCR in situ signal intensity within the TG (white dashed circle, Figure 1A-B) and found that mean *p2rx3b* intensity per µm³ was significantly reduced in *prdm16-/-* embryos (Figure 5D, n=5 per group), indicating decreased *p2rx3b* expression at the cellular level in *prdm16-/-* TG. Together, these findings suggest that *prdm16* plays a role in the differentiation process of the *p2rx3b*+ trigeminal sensory neurons from cNCCs.

**Figure 5.**
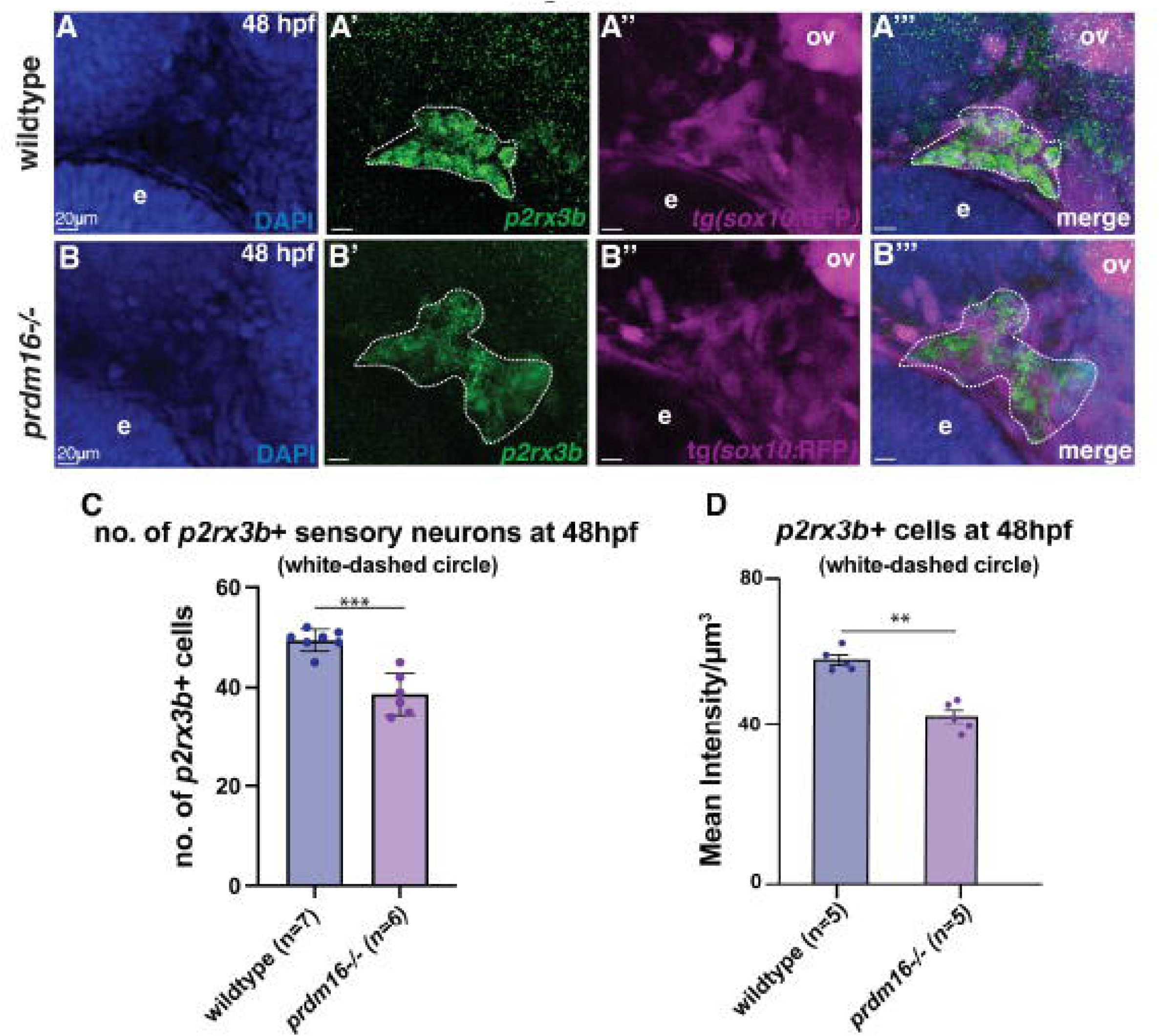
*prdm16* loss reduces *p2rx3b*+ expressing sensory neuron subtype differentiation in trigeminal ganglia at 48 hpf. Whole-mount HCR in situ hybridization was done for *p2rx3b* in Tg(*sox10*:RFP) background for wildtype and *prdm16*-/- embryos at 48 hpf. Lateral view, anterior to the left. Representative images show wildtype (A-A’’’) and *prdm16*−/− (C-C’’’) embryos at 48 hpf. DAPI (blue) marks nuclei, *p2rx3b* signal is shown in green, and *sox10*:RFP labels neural crest cells shown in magenta; followed by merged images. The trigeminal ganglion region is outlined with a white dashed line. (C) Quantification of the number of *p2rx3b*+ sensory neurons within the outlined TG at 48 hpf showed a significant reduction of *p2rx3b* expressing cells in TG in *prdm16*-/-compared to wildtype (wildtype n=7, *prdm16*−/− n=6, p < 0.001, ***). (D) Quantification of mean *p2rx3b* signal intensity per µm³ within the outlined white dashed line TG at 48 hpf using the Kruskal-Wallis non-parametric ANOVA statistical test (wildtype n=5, *prdm16*−/− n=5, p < 0.01, **). Scale bars: 20 µm. Abbreviations: TG, trigeminal ganglion; ov, otic vesicle; e, eye; hpf, hours post fertilization.

### *prdm16* is necessary to maintain proper neurogenesis transcriptional programs in *sox10+* cranial neural crest cells

To dissect the molecular mechanism of *prdm16’s* control of cNCC neuronal derivative differentiation, we analyzed the transcriptome of zebrafish cranial NCCs. We examined our previously published RNA-seq dataset from isolated *sox10*:EGFP-positive cranial NCCs from wildtype and *prdm16-/-* embryos at 48 hpf^52^. Using a Tg(*sox10:*EGFP) transgenic line that labels the neural crest in trigeminal ganglia, dorsal root ganglia and pharyngeal arches, fluorescence-activated cell sorting (FACS) followed by RNA sequencing (RNA-seq) was performed. Previously, we examined the chondrogenesis and Wnt-related gene expression in the previous publication ^52^, however, the results indicated that the cell fate for neuronal progenitor cells was also altered in *prdm16* mutants. To determine if neuronal-related transcripts were differentially misexpressed in the developing TG, we replotted the data as a volcano plot and heatmap to show the fold change of neuronal genes from the RNA-seq performed at 48 hpf (Figure 6A-B). We found that important transcription factors responsible for the complete neurogenesis process in TG, for example, *neurog1*, *neurod6a/b*, and *neurod4,* are downregulated in *prdm16-/-* compared to wildtype (marked in yellow dotted rectangle, Figure 6A-B). Confirmation of expression by HCR in situ hybridization shows *neurog1* and *neurod1* are expressed in the wildtype craniosensory ganglia region and are also highly expressed in the pharyngeal arches at 48 hpf, which is significantly reduced in *prdm16−/−* mutants (Figure 6C-D and Figure 6F-G). Quantification of the mean intensity of *neurog1* and *neurod1* HCR signals per μm^3^ showed a reduction in *neurog1* and *neurod1* signaling near the TG and pharyngeal arches (Figure 6E and Figure 6H). While reduction of *neurog1* expression is seen in all arches in *prdm16-/-*, *neurod1* is only reduced in pharyngeal arch 1. This confirms that there is a decrease in the neurogenesis program with the loss of *prdm16*. Interestingly, expression of important sensory neuron cell type-specific genes *trk2a/2b* and *trk3a/3b*, were also downregulated in *prdm16-/-* compared to wildtype (Figure 6A-B). Interestingly, we did not detect a significant change in *p2rx3b* expression in *prdm16-/-* embryos relative to wildtype expression. This could also be a technical zero counted as dropouts, since a few other subpopulations, like *p2rx2* or *trpa1a/b,* fold change values were also zero due to lower expression and did not pass the cut-off value set for fold change.

**Figure 6.**
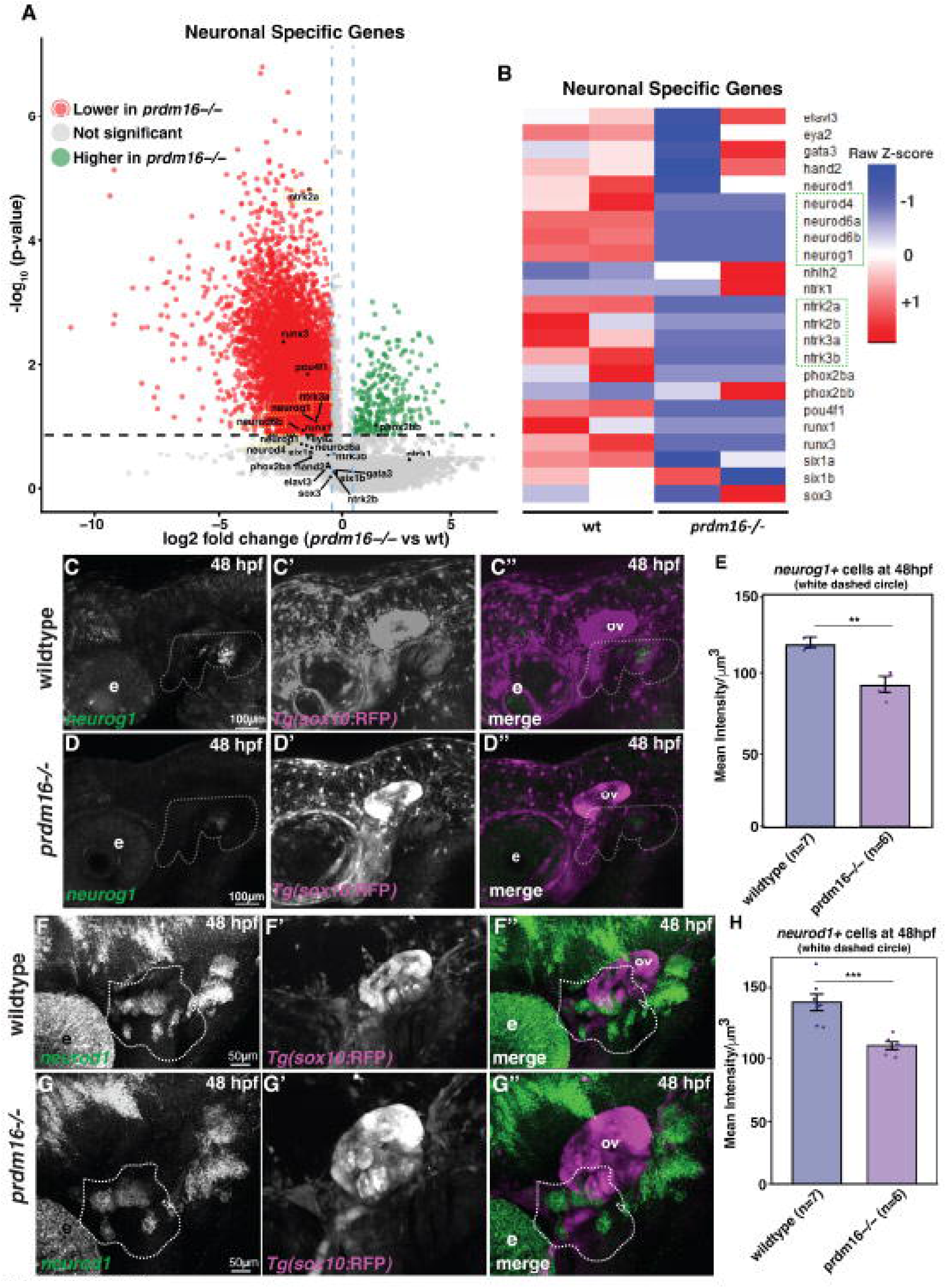
*prdm16* is required for the neurogenic transcriptional program in *sox10*+ cranial neural crest cells at 48 hpf. RNA-seq and HCR in situ hybridization show that loss of *prdm16* reduces expression of key neurogenic transcription factors and sensory-lineage markers in cranial neural crest derivatives. (A) Volcano plot of neuronal-associated transcripts from RNA-seq of FACS-isolated *sox10*:EGFP+ cranial neural crest cells at 48 hpf (*prdm16*-/- vs wildtype). Points are colored by differential expression category (red, downregulated in *prdm16*-/-; gray, not significant; green, upregulated in *prdm16*-/-). Dashed gray line indicates the fold-change cutoff used for visualization; yellow dashed boxes represent neurogenic (*neurog1*, *neurod6b*, *neurod1*) and sensory-lineage genes (*nrtk3a*, *ntrk2a*) that are downregulated in *prdm16*-/- embryos at 2dpf. (B) Heatmap showing row Z-scored expression of selected neuronal and sensory genes across wildtype and *prdm16*-/- samples, highlighting broad reduction of neurogenic transcription factors (e.g., *neurog1*, *neurod* family) and sensory-lineage markers (*trk* genes) in *prdm16*-/-. (C-D) Whole-mount HCR, lateral views anterior to the left for *neurog1* (green) together with *sox10*:RFP (magenta) in wildtype (C-C’’) and *prdm16*-/-(D-D’’) embryos at 48 hpf shows that, visually, there is a clear reduction in *neurog1*, neurogenesis-related transcription factor expression in *prdm16*-/- embryos compared to wildtype. The white dashed outline marks the cranial sensory ganglia and pharyngeal arch proximal region used for quantification. (E) Quantification of mean *neurog1* HCR signal intensity per µm³ within the dashed ROI at 48 hpf, showing moderately significant reduced signal in *prdm16*-/-(wildtype n=7; *prdm16*−/− n=6; p < 0.01, **). (F-G) Whole-mount HCR RNA-FISH for another important transcription factor for neurogenesis in TG, *neurod1* (green), together with *sox10*:RFP (magenta), showed a reduction of neurogenesis relate transcription factor*neurod1* HCR signal in *prdm16*-/- (G-G’’) compared to wildtype (F-F’’) embryos at 48 hpf. The dashed outline indicates the quantified region for the mean intensity of *neurod1* signal near the OV. (H) Quantification of mean *neurod1* HCR signal intensity per µm³ within the dashed ROI at 48 hpf, showing a significant reduction in *prdm16*-/- embryos (wildtype n=7; *prdm16*−/− n=6; p < 0.01, **). Scale bar: 100 µm (C-D), 50 µm (F-G). Abbreviations: ov, otic vesicle; e, eye; hpf, hours post fertilization.

### Loss of *Prdm16* in mice results in reduced TG size and neuron cell count

Finally, to determine if the function of PRDM16 in TG development is conserved across vertebrates, we examined the expression of *Prdm16* in mouse at embryonic day (E) 18.5, the latest stage of TG embryonic development. In E18.5 sections, PRDM16 co-immunostaining with HuC within the TG showed overlapping of HuC+ neuronal signal with PRDM16 (Figure 7A-A’’).

**Figure 7.**
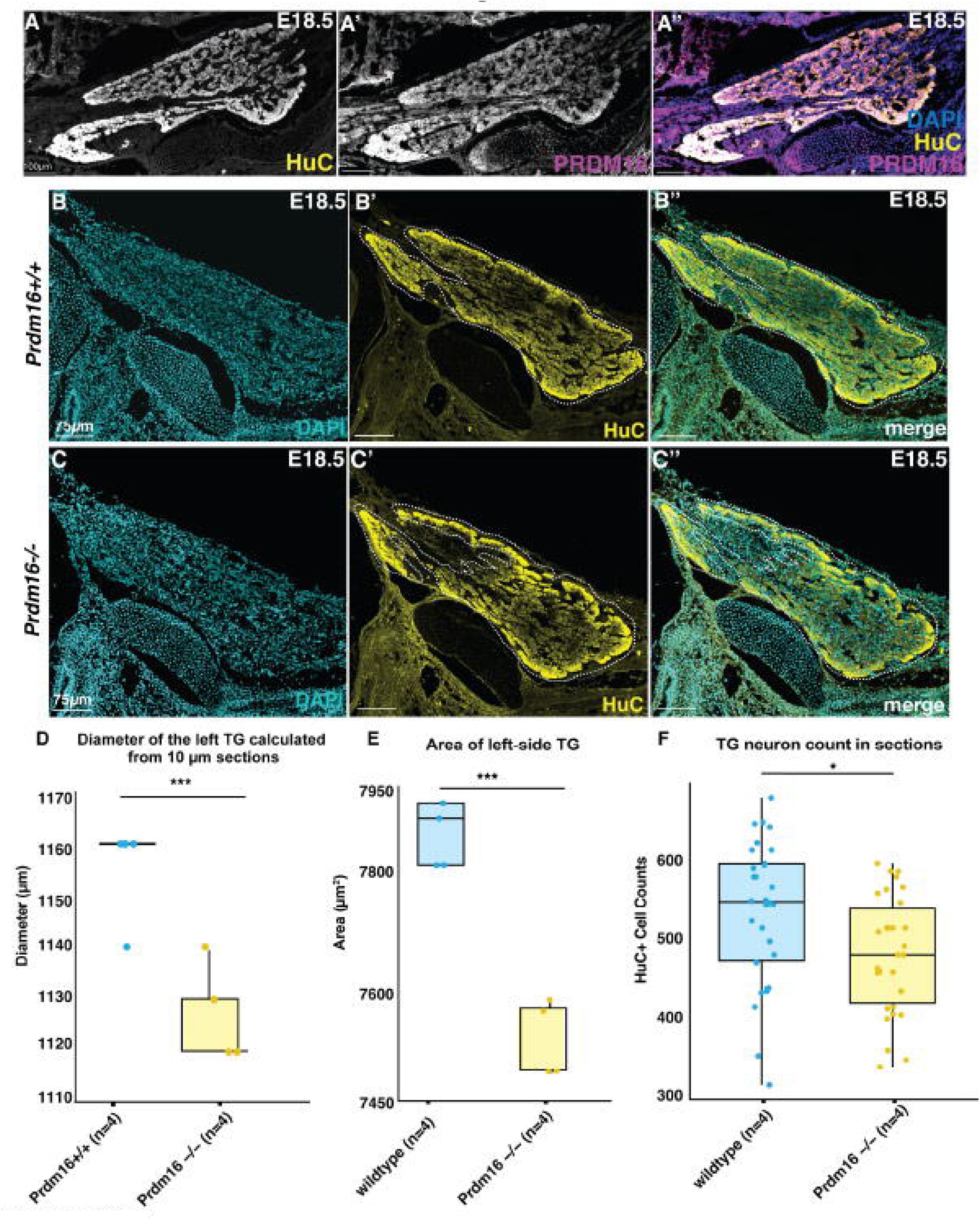
PRDM16 is expressed in late-gestation (E18.5) mouse trigeminal ganglion neurons, and *Prdm16^csp1^* mutants show reduced TG size and neuronal count. (A-A’’) Representative sagittal section (10 µm) through the E18.5 trigeminal ganglion (TG) immunostained for the pan-neuronal marker HuC (A, yellow in A’’) and PRDM16 (A’, magenta in A’’), with DAPI (blue) counterstain; merged image shows PRDM16 signal within the HuC+ TG domain. (B-C) Representative 10 µm sagittal sections of the left TG for wildtype, *Prdm16*^csp1/+^ (B-B’’) and *Prdm16^csp1/csp1^*mutant embryos at E18.5 stained for HuC (yellow) and DAPI (cyan); dashed outlines mark the TG region used for morphometric measurements in D-F. (D) TG span or diameter in dorsal-ventral extent quantified from serial 10 µm sections spanning the left TG (wildtype n=5, *Prdm16*+/− n=2, *Prdm16*−/− n=5). (E) Left TG area (µm²) measured at a representative mid-ganglion section (wildtype n=5, *Prdm16*^csp1/csp1^ n=5). (F) HuC+ neuron counts from anatomically matched TG sections (wildtype n=4, *Prdm16*^csp1/csp1^ n=4). Scale bars: 100 µm (A), 75 µm (B–C). p < 0.001, ***; p < 0.01, **; p <0.05, using the Kruskal-Wallis non-parametric ANOVA test.

Next, we examined TG development in the *Prdm16*^csp1^ mouse model. *Prdm16*^csp1^ homozygous mutant embryos display severe craniofacial defects at late gestation stages, including cleft palate and an open-mouth phenotype ^49^. We examined dissected head sagittal cryosections at E18.5 and used immunofluorescence to visualize TG neurons. We therefore asked if *Prdm16*^csp1/csp1^ embryos show altered TG size and neuronal content. HuC staining revealed well-defined TG in control embryos (Figure 7B-B’’), whereas the TG in *Prdm16* ^csp1/csp1^ embryos appeared smaller at comparable section levels (Figure 7C-C’’). We measured TG diameter or extent using the total number of serial 10 µm sections spanning the TG, and this value was significantly reduced in *Prdm16*^csp1^ mutants compared to wildtype mice at E18.5 (wildtype n=4, *Prdm16* ^csp1/+^ n=2, *Prdm16* ^csp1/csp1^ n=4, Figure 7D). We then measured TG area from a representative “mid-ganglion” plane by measuring the maximal TG diameter and multiplying by the corresponding height. Area (µm^2^) of the left side TG was significantly decreased in mutants compared to wildtype (n=4 for *Prdm16* ^+/+^ and *Prdm16* ^csp1/csp1^, Figure 7E). Finally, we counted HuC+ neurons in anatomically matched sections selected at consistent positions during sectioning, and *Prdm16* ^csp1/csp1^ showed a decrease in HuC+ cell numbers (n=4, p < 0.05) compared to wildtype (Figure 7F). Taken together, these results show that *Prdm16* loss is associated with a smaller TG volume and fewer TG neurons in mice, consistent with the reduced ganglion growth and neuronal accrual observed in zebrafish *prdm16* mutants.

## Discussion

In this work, we investigated the function of *prdm16* in TG development in both zebrafish and mouse. We show that *prdm16* is expressed and functions to promote the differentiation of TG neurons by controlling expression of downstream neuronal genes. Together with previous data, this supports the model that *prdm16* activates both the neuronal and cartilage GRN to promote differentiation of cNCC derivatives in the head. Classically thought of as a transcriptional repressor, there is evidence that *prdm16* activates transcriptional programs for neural crest specification and derivatives including craniofacial cartilage^52^.

In vertebrates, the trigeminal ganglion (TG) and other cranial ganglia have a dual origin where they arise from both cNCCs and neurogenic placodes^64,65^. Fate mapping analysis suggests that the proximal TG is neural crest derived while the distal portion of the ganglion are placode derived, although this relationship remains unresolved in zebrafish ^64,66,67^. Zebrafish TG sensory ganglia do develop as paired structures on either side of the head, positioned between the eye and the ear (Figure 1). The first trigeminal sensory neurons are born at around 11 hours post fertilization (hpf) and rapidly coalesce into a ganglion. By 24 hpf, the ganglia mediate the response to mechanical stimuli (touch) and chemical irritants^68,69^, resulting in a highly stereotypic escape behavior. In this context, using expression analysis, *prdm16* is expressed at the precise spatiotemporal window to influence TG morphogenesis. TG-proximal expression of *prdm16* emerges by approximately 18 hpf, persists through maturation, and is detectable in both neural crest–associated and placode-associated trigeminal cells during the period when these lineages coordinate to assemble a properly organized sensory ganglion. The contribution to the TG from both neural crest and placodes and how *prdm16* contributes to the assembly of the TG would be interesting to determine. The current data suggest that Prdm16 is expressed in a spatial and temporal pattern consistent with playing a role as a tissue-specific transcriptional regulator that contributes to coordinated TG development and sensory neuron differentiation.

Importantly, zebrafish *prdm16* loss results in a persistent reduction in TG neuron number and ganglion size, with decreased HuC+ neurons and reduced TG volume. There are several ways in which *prdm16*, acting as tissue specific transcription factor, could function to regulate cNCC development. One possibility is that *prmd16* controls cNCC specification and migration of cells to the ganglion itself. We tested this by examining the numbers and migration of cNCCs with live imaging in Tg(*sox10*:mRFP; *elavl3*:GFP) embryos. These data showed that the smaller size of TG did not result from a reduced total number of *sox10*+ neural crest cells reaching the TG region in *prdm16* mutants. This suggests that the differentiation defect arises following cNCC migration. Another mechanism is the regulation of cell proliferation and cell death within and around the developing TG. We examined the TG and did not observe a significant difference between wildtype and *prdm16-/-* embryos. This is consistent with previous results examining specifically the pharyngeal arch region w no changes were observed^51^. Thus, the most likely function of *prdm16* is to promote neuronal differentiation as demonstrated by fewer HuC+ neurons and fewer trigeminal projections and disrupted projection architecture, including reduced fine branching and abnormal local accumulations near the TG. This indicates the role of *prdm16* is as a regulator for the proper differentiation of cNCC-derived TG neurons and indicates that *prdm16* plays a role in proper neuronal cell fate differentiation from more restricted cNCC progenitor.

*prdm16*, along with its close paralogs, *prdm1a* and *prdm3,* have been shown to play a role in regulating the GRN for cell fate determination and differentiation in multiple development models ^45,70–72^. *prdm1a* (Blimp1) plays an important role in hematopoiesis, photoreceptor, muscle, limb and neural crest cell fate. Neural crest and Rohon-Beard sensory neurons are reduced in *prdm1a*-/- embryos in zebrafish and *prdm1a* plays a role in transcriptionally activating neural crest targets^73^. Interesting new work suggests *prdm1a* is an important cell fate switch between hair cells and lateral line mechanosensory cells^70^. Prdm3 and Prdm16 similarly have a roles in cell fate decisions including in lung^71^, cortical^74^, muscle/brown fat^72,75^ and craniofacial development^76^. In some contexts, Prdm3 and Prdm16 often act as a cell fate switch. Interesting, while very similar in protein structure, *prdm3* acts primarily as a repressor and *prdm16* acts as an activator, in craniofacial development ^52^. This role is consistent with the role we identify here with *prdm16* promoting the cell fate and differentiation of TG neurons. How it does so and what other fate might be affected in the ganglion as well as the function of its paralog *prdm3* in this process will be the subject of future study.

A key mechanistic insight from our study is that *prdm16* supports sensory neurogenesis programs and subtype differentiation. At the subtype level, *prdm16* mutants show a specific deficit in *p2rx3b*+ sensory neurons and reduced *p2rx3b* signal, indicating impaired differentiation of a defined nociceptor-associated population. In parallel, transcriptomic analysis of FACS-isolated *sox10*+ cranial neural crest cells revealed downregulation of core neurogenic transcription factors (including Neurog and Neurod family members) and sensory lineage-associated genes (e.g., trk receptors). Together, these results support a model in which *prdm16* regulates the GRN to promote a neurogenic state within cNCC–derived progenitors. *prdm16* normally may shift the balance of cNCC derivatives to promote neuronal differentiation, as well as cartilage differentiation and away from an alternate to be determined cell fate.

Finally, the mouse *Prdm16^csp1^* model supports an evolutionary conserved role of *Prdm16* in proper TG formation. PRDM16 protein is detectable in HuC+ TG neurons, indicating that PRDM16 is positioned to influence neuronal compartments of the ganglion in mouse embryos, similar to zebrafish. In *Prdm16^csp1^* homozygous mutant embryos, the TG is reduced in overall extent, decreased in estimated area, and contains fewer HuC+ neurons at matched anatomical levels. Although the severe late-gestation cleft palate defects in *Prdm16^csp1^*mutants could secondarily influence TG size^49^, the fact that we observe the same outcomes of reduced ganglion size and fewer HuC+ neurons in zebrafish supports a direct and conserved requirement for Prdm16 in TG development. Together, these data place Prdm16 as a transcriptional regulator that promotes trigeminal ganglion growth, sensory neurogenesis, and the maturation of trigeminal wiring patterns during cranial ganglion assembly.

## Materials and Methods

### Animals

#### Zebrafish

Zebrafish were housed and cared for using standard husbandry practices as described previously. Embryos were collected and raised in defined embryo water at 28.5°C, and developmental stages were assigned using published staging criteria^77^. The wild-type strain used in this study was AB. Transgenic lines included Tg(*sox10*:RFP)^78^ and Tg(*elavl3*:EGFP)^79^. These lines were crossed into the *prdm16* mutant backgrounds for experiments. The zebrafish *prdm16* mutant line was generated previously by CRISPR mutagenesis, similar to the allele described in *Shull et al*., 2020 ^51^. The allele used here (*prdm16^CO1027^* is predicted to contain a frameshift that disrupts the coding sequence upstream of the PR/SET domain, which is required for histone methyltransferase activity ^51,52^. All experiments were performed on embryos or larvae prior to sex determination (before 20 dpf). All zebrafish procedures were approved by the Institutional Animal Care and Use committee at the University of Minnesota, Twin Cities, and followed NIH guidelines for animal care.

#### Mice

*Prdm16^csp1^* ^49^ mice were obtained from The Jackson Laboratory and maintained on the FVB/NJ background and housed at 21-23° temperature under 12 h light/dark cycle with water and food provided ad libitum. For timed matings, *Prdm16^csp1/+^* heterozygous pairs were bred and the day a vaginal plug was detected in the morning was considered E0.5. Mice were euthanized by carbon dioxide inhalation and the embryos collected. We staged matched examined embryos at E18.5 from the same litter. Mice are bred and maintained according to a protocol approved by the Institutional Animal Care and Use committee at the University of Minnesota, Twin Cities, and followed NIH guidelines for animal care.

### Genotyping

#### Zebrafish

Fin clips, single embryos, or single larval tails were processed for genotyping. Samples were incubated in lysis buffer [10 mM Tris-HCl (pH 8.0), 50 mM KCl, 0.3% Tween-20, 0.3% NP-40, 1 mM EDTA] for 10 min at 95°C. Lysates were then digested with 50 µg Proteinase K at 55°C for 2 h, followed by Proteinase K inactivation at 95°C for 10 min. For genotyping the following *prdm16* primers were used for two exon sites: for exon site 1 (F) 5’ - TAAGCAATTATGTGATGCCGTC-3’ and (R) 5’ - CTTTTCACAGTCTTTGCACTCG-3’; for exon site 2 (F) 5’–TACACAGCAGTGTCAAGCCTTT – 3’ and (R) 5’– ACAAAAGAAAAGCGCAACAAAT – 3’. PCR was preformed using GoTaq G2 Green Master Mix (Promega) to genotype adult fin clips and embryos after immunofluorescence, and the PCR products were resolved on a 5% agarose gel. Each wildtype PCR product is about 200 bp with a 5 bp insertion in homozygous mutants for exon site 1 and 5bp deletion for exon site 2 in mutant alleles.

After HCR, zebrafish embryos were genotyped using the same DNA extraction method, but PCR was performed in M buffer [2 mM MgCl, 13.7 mM Tris-HCl (pH 8.4), 68.4 mM KCl, 0.001% gelatin, 1.8 mg/mL protease-free BSA, 136 µM each dATP/CTP/GTP/TTP] using GoTaq Flexi (Promega) and *prdm16* primers mentioned above. Amplicons were resolved on a 4% agarose gel.

#### Mice

*Prdm16*^csp1^ mice embryos were genotyped from the embryonic tissue tail tip. Genomic DNA was prepared using the Phire Tissue Direct PCR kit (ThermoFisher, F170S). Briefly, a ∼0.5 mm tissue piece was incubated in 20 µL Dilution Buffer with 0.5 µL DNA Release reagent for 2–5 min at room temperature, followed by enzyme inactivation at 98°C for 2 min. Lysates were briefly centrifuged, transferred to a clean tube, and stored at −20°C until PCR. PCR was carried out in 10 µL reactions containing 5 µL 2× Phire PCR Master Mix, 0.5 µL of each 10 µM primer, 0.5 µL DNA lysate, and nuclease-free water to volume. For genotyping *Prdm16*^csp1^ mice, we used the following primers: (F) 5′-AAGAGAGGCCTGGCAGTAA - 3′ and (R) 5′- GTGATGAACCCGTCAGTGAA - 3′. PCR products were digested with HindIII (Promega) at 37°C for 1 h. Digested products were run on a 4% agarose gel. The wildtype allele produces a single band at 344 bp, the *Prdm16^csp1^*homozygous mutants produce 211 bp and ∼133 bp fragments; heterozygotes show all bands.

Additionally, during E18.5 head dissections and cryopreservation for sectioning, embryos were screened for the characteristic cleft palate phenotype reported for Prdm16^csp1^ homozygous mutants (mutants older than ∼E14.5 can be recognized by the presence of a cleft palate). Consistent with published characterization of the Prdm16^csp1^ line ^48,49^, homozygous mutants can be recognized by the presence of a cleft palate. with associated craniofacial abnormalities (including altered tongue position). The severe craniofacial phenotype (including open-mouth appearance) at E18.5 helped identify likely *Prdm16^csp1/csp1^*for confirmation by PCR.

### HCR In Situ Hybridization

HCR probe sets (*neurog1, neurod1, p2rx3b, six1b, elavl3*) were obtained from Molecular Instruments. Whole-mount HCR was performed according to the manufacturer’s guidelines (Choi et al., 2016; Choi et al., 2018) and as described previously (Truong et al., 2023). Embryos were fixed overnight at 4°C in 4% PFA, rinsed in PBS, then dehydrated and permeabilized with gradual MeOH dehydration and finally two 10-min washes in 100% MeOH at room temperature. Samples were stored in fresh MeOH for at least 24 h at −20°C. Embryos were rehydrated through a graded MeOH/PBST (PBST: 1X PBS with 0.1% Tween-20) series (75%, 50%, 25%, 0%). For digestion, embryos were treated with proteinase K (10 µg/mL) for 30 secs (14 hpf), 3 min (24 hpf), and 15 min (48 hpf), washed twice in PBST, refixed in 4% PFA for 20 min, and washed five times in PBST. Embryos were incubated overnight at 37°C in the probes diluted in probe hybridization buffer. Hairpins were denatured at 95°C for 90 s and allowed to cool before amplification, and embryos were similarly incubated in hairpins and amplification buffer overnight at RT. After completion of the HCR reaction, embryos were kept in PBS at 4°C, protected from light. Whole embryos were mounted in 0.1% low-melt agarose and imaged on a Leica Stellaris 5. Images were processed using Z-project in ImageJ.

### Live Imaging

For time-lapse imaging and tracking, neural crest cells and neuronal cell bodies (with emphasis on the TG) were visualized using Tg(*sox10:*mRFP) and Tg(*elavl3:*GFP), respectively. We crossed these two lines to make a Tg(*sox10:*mRFP*;elavl3:*GFP) line to simultaneously visualize the migration of sox10+ cells in the TG area and visualize TG neurons by *elavl3*. Embryos were raised until the development stage of interest and prepared for live imaging by anesthetizing with clove oil (0.04% V/V clove oil in ethanol) solution diluted in embryo water. Embryos were embedded in 0.4% low-melt agarose and imaged on a Leica Stellaris 5 equipped with an incubator chamber to maintain 28°C during long acquisitions. Imaging was performed every 5 min from 22 hpf to 30 hpf, keeping the TG region in view. At each time point, we collected total ∼100 µm z-stacks at 1 µm z-steps with 5-minute intervals for ∼12 h. Movies and images were processed in ImageJ. After imaging, embryos were removed from agarose and genotyped using the *prdm16* genotyping protocol described above.

### Immunostaining

#### Zebrafish Whole-mount

Embryos were collected at the stated stages and fixed in 4% PFA for 2 h at 4°C. Fixation was stopped by washing in 3× PBS (pH 7.3). Samples were then washed three times in 1× PBS with 1% Triton X-100 for 10 min each at room temperature. Embryos were blocked for 1 h at room temperature in blocking solution containing 10% normal goat serum and 1% bovine serum albumin in 1× PBS. Primary antibodies were diluted 1:200 in blocking solution and incubated with samples overnight at 4°C. Antibodies used were anti-acetylated β-tubulin (Sigma, T6793) and anti-HuC (Invitrogen, 16A11). Following primary incubation, embryos were washed thoroughly in PBS with 0.1% Triton X-100, then incubated with the appropriate secondary antibodies (Alexa Fluor 488-conjugated donkey anti-mouse IgG, Invitrogen, A32766TR, and Alexa Fluor 594-conjugated donkey anti-mouse IgG, Invitrogen, A21203) overnight at 4°C. Samples were washed again in 1× PBS with 0.1% Triton X-100, counterstained with DAPI in 1×PBS for 20 min at room temperature, briefly rinsed in PBS with 0.1% Triton X-100, and mounted in 0.1% agarose on glass-bottom dishes for imaging.

For anti-phosphohistone H3 (proliferation) and anti-cleaved caspase 3 (cell death), we used a modified protocol. Embryos were fixed in 4% PFA overnight at 4°C at 24 hpf, then dehydrated through a methanol series and held in 100% methanol for at least one overnight. After rehydration through the reverse methanol series, embryos were washed 3× for 10 min in PBS (pH 7.3) containing 0.1% Tween. Samples were equilibrated in 150 mM Tris (pH 9.5) for 5 min, then incubated in the Tris solution at 70°C for 20 min. After antigen retrieval, embryos were washed 3× for 10 min in PBS + 0.1% Tween and then 2× for 5 min in distilled water. Embryos were blocked at room temperature in blocking solution (10% normal goat serum and 1% bovine serum albumin in 1× PBS), then incubated in primary antibodies for 2 days at 4°C. Primary antibodies were anti phosphohistone H3 (Sigma, H0412) and anti-cleaved caspase 3 (Cell Signaling, 9661), diluted (1:200) in blocking solution. After washing in PBS with 0.1% Triton X-100, embryos were incubated in goat anti-rabbit 546 secondary antibodies (Invitrogen) for 1 day at 4°C. Samples were washed thoroughly, incubated in DAPI (1mg/ml) diluted in PBS for 15 min at RT, washed, and taken through a glycerol/PBS gradient before mounting on slides with 75% glycerol with 25% PBS. All samples were imaged on a Leica Stellaris 5 confocal microscope. Image processing was performed using LAS X and ImageJ.

#### Mice Tissue Preparation for Sections

Mouse embryos were collected at E18.5 and rinsed in cold PBS. The heads of the animals were dissected, and the tails were saved for genotyping. Dissected embryo heads were fixed in 4% paraformaldehyde (PFA) in cold PBS at 4°C overnight with gentle rocking, then washed 3× in PBS (20 min each). For cryoprotection, embryos were incubated sequentially in 10% sucrose, 20% sucrose, and 30% sucrose in cold PBS, followed by 30% sucrose: OCT (1:1). Embryos were embedded in OCT in labeled cryomolds. Blocks were frozen on dry ice until solid and stored at −80°C until sectioning. For cryosectioning, OCT blocks were transferred from −80°C on dry ice and mounted onto a pre-cooled stub using OCT. Blocks were equilibrated in the cryostat chamber (−21 to −24°C) for ∼60 min before cutting. Sections were cut at 10 µm and collected on Superfrost Plus (ThermoFisher) charged slides. Slides were air-dried for 30 min and then held for at least 2 hr in 4°C prior to any PBS washes to minimize section loss.

#### Immunohistochemistry (IHC) on sections

Sections were washed 3× for 5 min in 1X PBS at room temperature. Heat-induced epitope retrieval (HIER) was performed using 10mM sodium citrate buffer (pH 6.0), first incubating 5min in RT and then in a cooker at 70°C with boiled 10mM sodium citrate buffer (pH 6.0), holding the slides with a rack. After HIER steps, slides were washed 2× for 5 min in PBS. Sections were blocked for 2 hr at room temperature in blocking solution (10% Donkey Serum in PBS-1% Triton-X solution + Fab fragments (40μg/ ml of AffiniPure® Fab Fragment Donkey Anti-Mouse IgG, Jackson Laboratories, product code: 715-007-003). Primary antibodies were diluted in the same blocking solution and applied at 100 µL per slide overnight at 4°C in a humidified slide chamber. Antibodies used were anti-HuC (Invitrogen, 16A11) and Prdm16 (BioTechne, AF6295). The dilutions for different antibodies are as follows: Prdm16 (1:200) and HuC (1:100). The next day, slides were washed 3× for 15 min in PBS with 0.1% Triton X-100 at room temperature. Fluorophore-conjugated secondary antibodies (Alexa Fluor 488-conjugated donkey anti-mouse IgG, Invitrogen, A32766TR, and Alexa Fluor 594-conjugated donkey anti-mouse IgG, Invitrogen, A21203) were diluted in blocking solution and incubated on sections (100 µL/slide) protected from light for 3 hr. After secondary incubation, slides were washed 3× for 15 min in PBS with 0.1% Triton X-100, then washed 2× for 5 min in PBS. Nuclei were counterstained for DAPI (1mg/ml) in 1×PBS (1:1000 dilution) incubation for 20 mins at RT. For autofluorescence reduction, 1X TrueBlack® Lipfuscin Autofluorescence reagent (Biotium, diluted in 70% ethanol) was used, followed by 2X PBS washes. Sections were dried, mounted with Vectashield Mounting Media (Vector Laboratories), coverslipped and sealed. Slides were kept in 4°C protected from light until imaging. For analysis of trigeminal ganglia on sections, comparable anatomical levels were identified across embryos, and images were collected using identical settings on a Leica Stellaris 5. TG size measurements and neuron counts were quantified from matched sections using consistent ROIs across genotypes. Images were quantified in ImageJ using the cell counter plug-in.

#### 3D quantification of HuC+ cells in IMARIS

To quantify the number of HuC+ TG neurons in 3D across all z-stacks and estimate the volume of TG of whole-mount zebrafish embryos, we used a licensed image analysis software, IMARIS 11.0 (https://imaris.oxinst.com/). To segment cells detecting both nuclei (stained with DAPI) and cell body (HuC marks neuronal cell body), we used the “Imaris Cell” package (license bought by University of Minnesota Imaging Centers, UIC). First, the Leica Stellaris 5 files (.lif) were converted to IMARIS compatible image files (.ims) using the software IMARIS file converter 11.0. The trigeminal ganglion at 48 hpf was cropped using the “crop 3D” feature available in IMARIS, which allows cropping images in xyz axes, which is important for 3D counting. The images were post-processed for both the DAPI channel and the HuC channel, selecting a median filter (3*3*3*) and γ-adjustment = 0.90 in the image process function in IMARIS. First, nuclei were detected using the signal for the DAPI channel: the nuclei diameter was set to 3.50 μm and nuclei were split by seed points, ensuring one nucleus per cell. Next, on the same analysis cell bodies were detected using the HuC channel signal, and the cell’s smallest diameter was set to 4.50 μm with a membrane detail of 0.450 μm. With these consistent parameters set, cell area, volume, and cell count were determined in 3D for both wild-type and *prdm16-/-* zebrafish embryos.

#### Bioinformatics and Statistical Analysis

Complete RNA-seq datasets are available at the Gene Expression Omnibus repository (GSE175767). After downloading the RNA-seq data, identified the neuron differentiation or neurogenesis-related genes using ‘GOBP_Neuron_Differentiation’ (https://amigo.geneontology.org/amigo/term/GO:0030182). We conducted differential expression analysis using the Bioconductor ‘DESeq2’ package in R (https://bioconductor.org/packages/release/bioc/html/DESeq2.html). Then we plotted the genes in volcano plot and heatmap to compare the neuron differentiation program among wildtype and *prdm16-/-* embryos at 48 hpf.

All the statistical analyses were done in GraphPad Prism (version 10.6.0) using the Kruskal-Wallis non-parametric ANOVA test. The p-values for each analysis are reported for the respective figure panels in the figure legends.

## Acknowledgements

We thank Julaine Roffers-Agarwal for help with mouse line maintenance and training, Bryan Zepeda for help with live cell imaging, past and present members of the Artinger lab for project feedback, Marc Tye at the University of Minnesota, and the zebrafish and mouse facility team for excellent animal care. This work is supported by NIDCR R01DE030629 to K.B.A.

---

**Supplementary Figure 1. *prdm16* is expressed in *six1b*+ placode-derived cells in the TG region at 48 hpf.** (A-A’’) Whole-mount HCR for *six1b* (yellow; placodal domain) and *prdm16* (magenta) in zebrafish embryos at 48 hpf. Panels show *six1b* alone (A), *prdm16* alone (A’’), and the merged image (A’’). The trigeminal ganglion (TG) region is outlined with a white dashed circle. Abbreviations: ov, otic vesicle; e, eye; hpf, hours post fertilization. Scale bar: 50 µm.

**Supplemental Figure 2.** T**G morphology is consistently reduced in *prdm16−/−* embryos across stages, but with variable severity.** (A-B) Confocal images of HuC immunofluorescence (green) with DAPI (blue) at 24 hpf, showing TG in wildtype (A–A’) and a smaller HuC+ TG domain in *prdm16−/−* embryos (B-B’). The TG is outlined (white dashed circle). Scale bar: 100 µm. (C-E) Confocal images at 48 hpf showing HuC (magenta), Tg(*sox10*:RFP) (yellow), and DAPI (blue). Wildtype embryos form a well-defined TG (C-C’’), whereas *prdm16*−/− embryos show a reduced TG with altered organization, including a commonly observed phenotype (D-D’’; n=11/16) and a more severe variation is shown (E-E’’; n=5/16). e, eye; ov, otic vesicle; are labeled for orientation. Scale bar: 50 µm.

**Supplementary Figure 3. No detectable change in proliferation or apoptosis in the TG or TG-proximal region at 24 hpf in *prdm16*−/− embryos.** (A-B) Whole-mount immunostaining for anti-phosphohistone H3 (pHH3, green; proliferating cells) in wildtype (A-A’’’) and *prdm16*-/- embryos (B-B’’’) at 24 hpf, shown with Tg(*sox10*:RFP) (magenta) and merged views. The TG and TG-proximal region between eye (e) and otic vesicle (ov), near pharyngeal arch region (pa) is outlined (white dashed line) and used for quantification. Scale bar: 50 µm. (C-D) Whole-mount immunostaining for anti-cleaved caspase-3 (CC3, green; apoptotic cells) in wildtype (C-C’’’) and *prdm16*-/- embryos (D-D’’’) at 24 hpf, shown with brightfield and merged views. The quantified region is outlined (white dashed line). Scale bar: 100 µm. (E-F) Quantification of pHH3+ cells (E) and cleaved caspase-3+ cells (F) within the outlined region shows no significant difference between genotypes (n=4 per group; ns in the Kruskal-Wallis non-parametric ANOVA test).

**Supplemental Movie 1.** Time-lapse imaging of a Tg(*sox10*:mRFP; *elavl3*:GFP) wildtype embryo with *sox10*+ neural crest cells (cyan) are migrating to complete the formation of the trigeminal ganglia, while trigeminal sensory neurons are marked with *elavl3*, shown in yellow. Live Imaging was started at 22 hpf (0 hr) when the TG is starting to coalesce and followed embryos for 13 hours (to approximately 35 hpf). Scale bar: 50 µm.

**Supplemental Movie 2.** Time-lapse imaging of a Tg(*sox10*:mRFP; *elavl3*:GFP) *prdm16*-/- embryo showing the same number of *sox10*+ neural crest cells (cyan) are migrating to complete the formation of the trigeminal ganglia compared to wildtype from supplementary video 1, while trigeminal sensory neurons are marked with elavl3, shown in yellow. The TG in *prdm16*-/- looks smaller compared to the wildtype shown in supplementary video 1 across the time-lapse video. Live Imaging was started at 22 hpf (0 hr) when the TG is starting to coalesce, and followed embryos for 13 hours (to approximately 35 hpf). Scale bar: 50 µm.

## References

1. Northcutt RG, Gans C. The Genesis of Neural Crest and Epidermal Placodes: A Reinterpretation of Vertebrate Origins. The Quarterly Review of Biology. 1983;58(1):1–28. 10.1086/413055

2. Streit A. The cranial sensory nervous system: specification of sensory progenitors and placodes. StemBook. Published online 2008. 10.3824/stembook.1.31.1

3. Mishina Y, Snider TN. Neural crest cell signaling pathways critical to cranial bone development and pathology. Experimental Cell Research. 2014;325(2):138–147. 10.1016/j.yexcr.2014.01.019

4. Dash S, Trainor PA. The development, patterning and evolution of neural crest cell differentiation into cartilage and bone. Bone. 2020;137:115409. 10.1016/j.bone.2020.115409

5. Mizuseki K, Sakamoto T, Watanabe K, et al. Generation of neural crest-derived peripheral neurons and floor plate cells from mouse and primate embryonic stem cells. Proc Natl Acad Sci USA. 2003;100(10):5828–5833. 10.1073/pnas.1037282100

6. Newbern JM. Molecular Control of the Neural Crest and Peripheral Nervous System Development. In: Current Topics in Developmental Biology. Vol 111. Elsevier; 2015:201–231. 10.1016/bs.ctdb.2014.11.007

7. Méndez-Maldonado K, Vega-López GA, Aybar MJ, Velasco I. Neurogenesis From Neural Crest Cells: Molecular Mechanisms in the Formation of Cranial Nerves and Ganglia. Front Cell Dev Biol. 2020;8:635. 10.3389/fcell.2020.00635

8. Umehara Y, Toyama S, Tominaga M, et al. Robust induction of neural crest cells to derive peripheral sensory neurons from human induced pluripotent stem cells. Sci Rep. 2020;10(1):4360. 10.1038/s41598-020-60036-z

9. Kalcheim C. The association between neural crest-derived glia and melanocyte lineages throughout development and disease. Developmental Dynamics. Published online November 11, 2025:dvdy.70098. 10.1002/dvdy.70098

10. Sauka-Spengler T, Bronner-Fraser M. A gene regulatory network orchestrates neural crest formation. Nat Rev Mol Cell Biol. 2008;9(7):557–568. 10.1038/nrm2428

11. Simões-Costa M, Bronner ME. Establishing neural crest identity: a gene regulatory recipe. Development. 2015;142(2):242–257. 10.1242/dev.105445

12. Hovland AS, Rothstein M, Simoes-Costa M. Network architecture and regulatory logic in neural crest development. WIREs Mechanisms of Disease. 2020;12(2):e1468. 10.1002/wsbm.1468

13. Soldatov R, Kaucka M, Kastriti ME, et al. Spatiotemporal structure of cell fate decisions in murine neural crest. Science. 2019;364(6444):eaas9536. 10.1126/science.aas9536

14. Bi-Lin KW, Seshachalam PV, Tuoc T, Stoykova A, Ghosh S, Singh MK. Critical role of the BAF chromatin remodeling complex during murine neural crest development. Conlon FL, ed. PLoS Genet. 2021;17(3):e1009446. 10.1371/journal.pgen.1009446

15. Fountain DM, Sauka-Spengler T. The SWI/SNF Complex in Neural Crest Cell Development and Disease. Annu Rev Genom Hum Genet. 2023;24(1):203–223. 10.1146/annurev-genom-011723-082913

16. Barlow LA. Cranial Nerve Development: Placodal Neurons Ride the Crest. Current Biology. 2002;12(5):R171–R173. 10.1016/S0960-9822(02)00734-0

17. Halmi CA, Wu CY, Taneyhill LA. Neural crest cell-placodal neuron interactions are mediated by Cadherin-7 and N-cadherin during early chick trigeminal ganglion assembly. F1000Res. 2022;11:741. 10.12688/f1000research.122686.2

18. Barlow LA. Cranial Nerve Development: Placodal Neurons Ride the Crest. Current Biology. 2002;12(5):R171–R173. 10.1016/S0960-9822(02)00734-0

19. Leonard CE, McIntosh A, Sanyal J, Taneyhill LA. The transcriptional landscape of the developing chick trigeminal ganglion. Developmental Biology. 2025;520:108–116. 10.1016/j.ydbio.2024.12.013

20. Rocha M, Singh N, Ahsan K, Beiriger A, Prince VE. Neural crest development: insights from the zebrafish. Developmental Dynamics. 2020;249(1):88–111. 10.1002/dvdy.122

21. Graham A. The neural crest. Current Biology. 2003;13(10):R381–R384. 10.1016/S0960-9822(03)00315-4

22. Graham A, Begbie J, McGonnell I. Significance of the cranial neural crest. Developmental Dynamics. 2004;229(1):5–13. 10.1002/dvdy.10442

23. Köntges G, Lumsden A. Rhombencephalic neural crest segmentation is preserved throughout craniofacial ontogeny. Development. 1996;122(10):3229–3242. 10.1242/dev.122.10.3229

24. Minoux M, Rijli FM. Molecular mechanisms of cranial neural crest cell migration and patterning in craniofacial development. Development. 2010;137(16):2605–2621. 10.1242/dev.040048

25. Jia S, Liu J, Chu Y, Liu Q, Mai L, Fan W. Single-cell RNA sequencing reveals distinct transcriptional features of the purinergic signaling in mouse trigeminal ganglion. Front Mol Neurosci. 2022;15:1038539. 10.3389/fnmol.2022.1038539

26. Nguyen MQ, Wu Y, Bonilla LS, Von Buchholtz LJ, Ryba NJP. Diversity amongst trigeminal neurons revealed by high throughput single cell sequencing. Obukhov AG, ed. PLoS ONE. 2017;12(9):e0185543. 10.1371/journal.pone.0185543

27. Woo SH, Lukacs V, De Nooij JC, et al. Piezo2 is the principal mechanotransduction channel for proprioception. Nat Neurosci. 2015;18(12):1756–1762. 10.1038/nn.4162

28. Hasegawa H, Wang F. Visualizing mechanosensory endings of TrkC-expressing neurons in HS3ST-2-hPLAP mice. J of Comparative Neurology. 2008;511(4):543–556. 10.1002/cne.21862

29. Salas MM, Hargreaves KM, Akopian AN. TRPA1-mediated responses in trigeminal sensory neurons: interaction between TRPA1 and TRPV1. Eur J of Neuroscience. 2009;29(8):1568–1578. 10.1111/j.1460-9568.2009.06702.x

30. Mishra SK, Tisel SM, Orestes P, Bhangoo SK, Hoon MA. TRPV1-lineage neurons are required for thermal sensation. EMBO J. 2011;30(3):582–593. 10.1038/emboj.2010.325

31. Burnstock G. P2X receptors in sensory neurones. British Journal of Anaesthesia. 2000;84(4):476–488. 10.1093/oxfordjournals.bja.a013473

32. Norton WHJ, Rohr KB, Burnstock G. Embryonic expression of a P2X 3 receptor encoding gene in zebrafish. Mechanisms of Development. 2000;99(1-2):149–152. 10.1016/S0925-4773(00)00472-X

33. Li M, Wang Y, Banerjee R, et al. Molecular mechanisms of human P2X3 receptor channel activation and modulation by divalent cation bound ATP. eLife. 2019;8:e47060. 10.7554/eLife.47060

34. Fabbretti E. ATP P2X3 receptors and neuronal sensitization. Front Cell Neurosci. 2013;7. 10.3389/fncel.2013.00236

35. Koizumi M, Asano S, Furukawa A, et al. P2X3 receptor upregulation in trigeminal ganglion neurons through TNFα production in macrophages contributes to trigeminal neuropathic pain in rats. J Headache Pain. 2021;22(1):31. 10.1186/s10194-021-01244-4

36. Chen Y, Chen L, Ji T, Yu Y, Zhang T, Wang L. The purinergic receptor P2X3 promotes facial pain by activating neurons and cytokines in the trigeminal ganglion. International Immunopharmacology. 2024;130:111801. 10.1016/j.intimp.2024.111801

37. Mah W, Lee SM, Lee J, et al. A role for the purinergic receptor P2X3 in astrocytes in the mechanism of craniofacial neuropathic pain. Sci Rep. 2017;7(1):13627. 10.1038/s41598-017-13561-3

38. Erzurumlu RS, Murakami Y, Rijli FM. Mapping the face in the somatosensory brainstem. Nat Rev Neurosci. 2010;11(4):252–263. 10.1038/nrn2804

39. Özdinler PH, Erzurumlu RS. Slit2, a Branching–Arborization Factor for Sensory Axons in the Mammalian CNS. J Neurosci. 2002;22(11):4540–4549. 10.1523/JNEUROSCI.22-11-04540.2002

40. Ulupinar E, Datwani A, Behar O, Fujisawa H, Erzurumlu R. Role of Semaphorin III in the Developing Rodent Trigeminal System. Molecular and Cellular Neuroscience. 1999;13(4):281–292. 10.1006/mcne.1999.0747

41. Eng SR, Gratwick K, Rhee JM, Fedtsova N, Gan L, Turner EE. Defects in Sensory Axon Growth Precede Neuronal Death in Brn3a-Deficient Mice. J Neurosci. 2001;21(2):541–549. 10.1523/JNEUROSCI.21-02-00541.2001

42. Gatto G, Dudanova I, Suetterlin P, et al. Protein Tyrosine Phosphatase Receptor Type O Inhibits Trigeminal Axon Growth and Branching by Repressing TrkB and Ret Signaling. J Neurosci. 2013;33(12):5399–5410. 10.1523/JNEUROSCI.4707-12.2013

43. Ding H, Clouthier DE, Artinger KB. Redundant roles of PRDM family members in zebrafish craniofacial development. Developmental Dynamics. 2013;242(1):67–79. 10.1002/dvdy.23895

44. Horn KH, Warner DR, Pisano M, Greene RM. PRDM16 expression in the developing mouse embryo. Acta Histochemica. 2011;113(2):150–155. 10.1016/j.acthis.2009.09.006

45. Hohenauer T, Moore AW. The Prdm family: expanding roles in stem cells and development. Development. 2012;139(13):2267–2282. 10.1242/dev.070110

46. Pinheiro I, Margueron R, Shukeir N, et al. Prdm3 and Prdm16 are H3K9me1 Methyltransferases Required for Mammalian Heterochromatin Integrity. Cell. 2012;150(5):948–960. 10.1016/j.cell.2012.06.048

47. Zhou B, Wang J, Lee SY, et al. PRDM16 Suppresses MLL1r Leukemia via Intrinsic Histone Methyltransferase Activity. Molecular Cell. 2016;62(2):222–236. 10.1016/j.molcel.2016.03.010

48. Bjork BC, Gomez AC, Ahmed A, et al. Prdm16 and Mecom mutants exhibit cleft secondary palate as a result of perturbations that affect different stages of palatogenesis. The FASEB Journal. 2018;32(S1). 10.1096/fasebj.2018.32.1_supplement.776.7

49. Bjork BC, Turbe-Doan A, Prysak M, Herron BJ, Beier DR. Prdm16 is required for normal palatogenesis in mice. Human Molecular Genetics. 2010;19(5):774–789. 10.1093/hmg/ddp543

50. Strassman A, Schnütgen F, Dai Q, et al. Generation of a multipurpose *Prdm16* mouse allele by targeted gene trapping. Disease Models & Mechanisms. 2017;10(7):909–922. 10.1242/dmm.029561

51. Shull LC, Sen R, Menzel J, Goyama S, Kurokawa M, Artinger KB. The conserved and divergent roles of Prdm3 and Prdm16 in zebrafish and mouse craniofacial development. Developmental Biology. 2020;461(2):132–144. 10.1016/j.ydbio.2020.02.006

52. Shull LC, Lencer ES, Kim HM, et al. PRDM paralogs antagonistically balance Wnt/β-catenin activity during craniofacial chondrocyte differentiation. Development. 2022;149(4):dev200082. 10.1242/dev.200082

53. Chasman DI, Schürks M, Anttila V, et al. Genome-wide association study reveals three susceptibility loci for common migraine in the general population. Nat Genet. 2011;43(7):695–698. 10.1038/ng.856

54. Lee H, Chen C, Ong J, et al. Association of rs2651899 Polymorphism in the Positive Regulatory Domain 16 and Common Migraine Subtypes: A Meta-Analysis. Headache. 2020;60(1):71–80. 10.1111/head.13670

55. Antonellis A, Huynh JL, Lee-Lin SQ, et al. Identification of Neural Crest and Glial Enhancers at the Mouse Sox10 Locus through Transgenesis in Zebrafish. Barsh GS, ed. PLoS Genet. 2008;4(9):e1000174. 10.1371/journal.pgen.1000174

56. Wada N, Javidan Y, Nelson S, Carney TJ, Kelsh RN, Schilling TF. Hedgehog signaling is required for cranial neural crest morphogenesis and chondrogenesis at the midline in the zebrafish skull. Development. 2005;132(17):3977–3988. 10.1242/dev.01943

57. Nord H, Skalman LN, Von Hofsten J. Six1 regulates proliferation of Pax7+ muscle progenitors in zebrafish. Journal of Cell Science. Published online January 1, 2013:jcs.119917. 10.1242/jcs.119917

58. Van Der Sar AM, Živković D, Den Hertog J. Eye defects in receptor protein-tyrosine phosphatase α knock-down zebrafish. Developmental Dynamics. 2002;223(2):292–297. 10.1002/dvdy.10059

59. Sánchez E, Azcona LJ, Paisán-Ruiz C. Pla2g6 Deficiency in Zebrafish Leads to Dopaminergic Cell Death, Axonal Degeneration, Increased β-Synuclein Expression, and Defects in Brain Functions and Pathways. Mol Neurobiol. 2018;55(8):6734–6754. 10.1007/s12035-017-0846-2

60. Pan YA, Choy M, Prober DA, Schier AF. Robo2 determines subtype-specific axonal projections of trigeminal sensory neurons. Development. 2012;139(3):591–600. 10.1242/dev.076588

61. Gau P, Curtright A, Condon L, Raible DW, Dhaka A. An ancient neurotrophin receptor code; a single Runx/Cbfβ complex determines somatosensory neuron fate specification in zebrafish. Granato M, ed. PLoS Genet. 2017;13(7):e1006884. 10.1371/journal.pgen.1006884

62. Caron SJC, Prober D, Choy M, Schier AF. In vivo birthdating by BAPTISM reveals that trigeminal sensory neuron diversity depends on early neurogenesis. Development. 2008;135(19):3259–3269. 10.1242/dev.023200

63. Kucenas S, Soto F, Cox JA, Voigt MM. Selective labeling of central and peripheral sensory neurons in the developing zebrafish using P2X3 receptor subunit transgenes. Neuroscience. 2006;138(2):641–652. 10.1016/j.neuroscience.2005.11.058

64. Noden DM. Spatial integration among cells forming the cranial peripheral nervous system. J Neurobiol. 1993;24(2):248–261. 10.1002/neu.480240210

65. Covell DA, Noden DM. Embryonic development of the chick primary trigeminal sensory-motor complex. J of Comparative Neurology. 1989;286(4):488–503. 10.1002/cne.902860407

66. Stark MR. Vertebrate neurogenic placode development: Historical highlights that have shaped our current understanding. Developmental Dynamics. 2014;243(10):1167–1175. 10.1002/dvdy.24152

67. Hamburger V. Experimental analysis of the dual origin of the trigeminal ganglion in the chick embryo. J Exp Zool. 1961;148(2):91–123. 10.1002/jez.1401480202

68. Sagasti A, Guido MR, Raible DW, Schier AF. Repulsive Interactions Shape the Morphologies and Functional Arrangement of Zebrafish Peripheral Sensory Arbors. Current Biology. 2005;15(9):804–814. 10.1016/j.cub.2005.03.048

69. Brehm N, Wenke N, Glessner K, Haehnel-Taguchi M. Physiological responses of mechanosensory systems in the head of larval zebrafish (Danio rerio). Front Robot AI. 2023;10:1212626. 10.3389/frobt.2023.1212626

70. Sandler JE, Tsai YY, Chen S, et al. prdm1a drives a fate switch between hair cells of different mechanosensory organs. Nat Commun. 2025;16(1):7662. 10.1038/s41467-025-62942-0

71. He H, Bell SM, Davis AK, et al. PRDM3/16 regulate chromatin accessibility required for NKX2-1 mediated alveolar epithelial differentiation and function. Nat Commun. 2024;15(1):8112. 10.1038/s41467-024-52154-3

72. Seale P, Bjork B, Yang W, et al. PRDM16 controls a brown fat/skeletal muscle switch. Nature. 2008;454(7207):961–967. 10.1038/nature07182

73. Hernandez-Lagunas L, Choi IF, Kaji T, et al. Zebrafish narrowminded disrupts the transcription factor prdm1 and is required for neural crest and sensory neuron specification. Developmental Biology. 2005;278(2):347–357. 10.1016/j.ydbio.2004.11.014

74. He L, Wen J, Dai Q. PRDM16 functions as a co-repressor in the BMP pathway to suppress neural stem cell proliferation. eLife. 2025;14:RP104076. 10.7554/eLife.104076.3

75. Kajimura S, Seale P, Kubota K, et al. Initiation of myoblast to brown fat switch by a PRDM16–C/EBP-β transcriptional complex. Nature. 2009;460(7259):1154–1158. 10.1038/nature08262

76. Denipah-Cook Q, Saxton BV, Artinger KB, Shull LC. PRDM paralogs are required for Meckel’s cartilage formation during mandibular bone development. Developmental Biology. 2026;530:210–223. 10.1016/j.ydbio.2025.12.002

77. Kimmel CB, Ballard WW, Kimmel SR, Ullmann B, Schilling TF. Stages of embryonic development of the zebrafish. Developmental Dynamics. 1995;203(3):253–310. 10.1002/aja.1002030302

78. Blasky AJ, Pan L, Moens CB, Appel B. Pard3 regulates contact between neural crest cells and the timing of Schwann cell differentiation but is not essential for neural crest migration or myelination. Developmental Dynamics. 2014;243(12):1511–1523. 10.1002/dvdy.24172

79. Sato T, Takahoko M, Okamoto H. HuC:Kaede, a useful tool to label neural morphologies in networks in vivo. genesis. 2006;44(3):136–142. 10.1002/gene.20196

