## Supplementary figures and images for "PRDM16 is necessary for sensory neuronal development in the Trigeminal Ganglion"

### Supplemental figure1

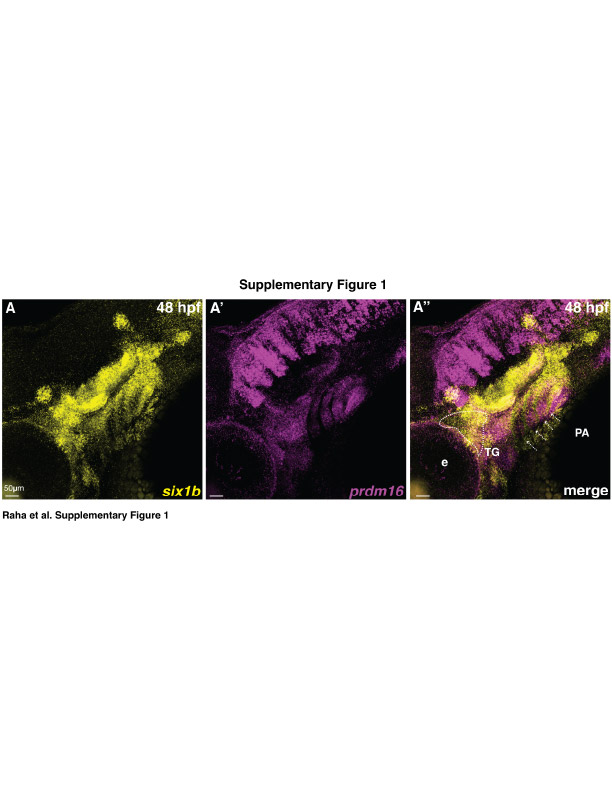

### Supplemental figure2

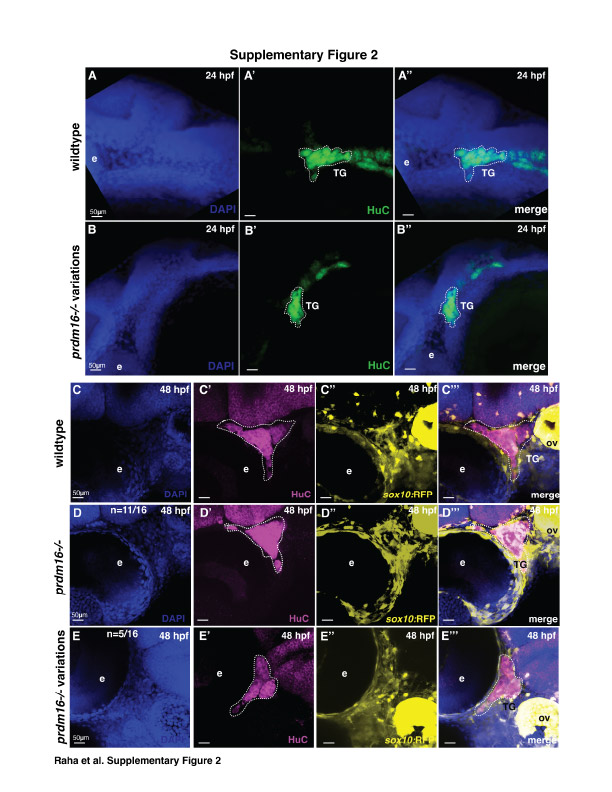

### Supplemental figure3

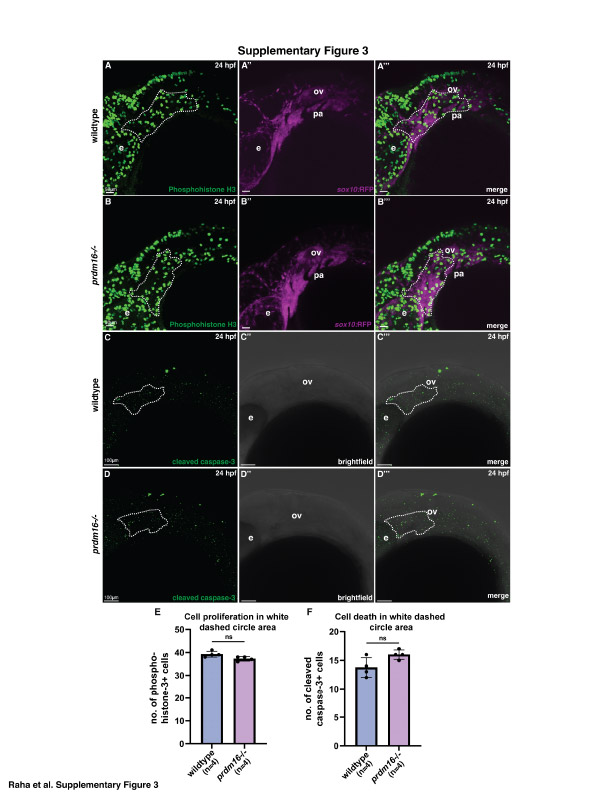
